# Tumor hypoxia adaptation depends on FLAD1 mediated mitochondrial metabolic reprogramming

**DOI:** 10.1101/2025.03.05.641576

**Authors:** Xiangyu Zhao, Tao Wu, Sanan Wu, Yujin Chen, Yu Zhang, Jing Chen, Kaixuan Diao, Zaoke He, Jiawei Yan, Tianzhu Lu, Chao Xu, Lu Liu, Gaofeng Fan, Dongliang Xu, Xinxiang Li, Xiaopeng Xiong, Jianjun Cheng, Fang Bai, Xue-Song Liu

**Affiliations:** School of Life Science and Technology, ShanghaiTech University, Shanghai 201203, China; NHC Key Laboratory of Personalized Diagnosis and Treatment of Nasopharyngeal Carcinoma (Jiangxi Cancer Hospital of Nanchang University), Nanchang, Jiangxi, P.R.China; Shanghai Institute for Advanced Immunochemical Studies and School of Life Science and Technology, ShanghaiTech University, Shanghai, 201210, China; Department of Urology, Shuguang Hospital, Shanghai University of Traditional Chinese Medicine, Shanghai 200120, China; Department of Colorectal Surgery, Fudan University Shanghai Cancer Center, 270 Dong’an Road, Xuhui, Shanghai, 200032, China; School of Life Science and Technology, ShanghaiTech University, Shanghai 201203, China; Shanghai Clinical Research and Trial Center, Shanghai, China

**Keywords:** Hypoxia, Deep learning, Metabolic dependency, FLAD1, Therapeutic target

## Abstract

Hypoxia, a hallmark of solid tumors, promotes the malignant progression and is challenging to target. Metabolic reprogramming and the resulting metabolic vulnerabilities provide a promising strategy to target tumor hypoxia. Here we systematically compared the metabolic network differences between hypoxic and non-hypoxic cells, and developed a deep learning model, “DepFormer”, to predict the dependent metabolic genes in hypoxic tumor cells. The performance of DepFormer was validated using CRISPR screening dataset. Oxidative phosphorylation was identified as the most significantly hypoxia-dependent metabolic pathway, and *FLAD1* was predicted to be one of the key hypoxia-dependent metabolic genes. *FLAD1* locus is amplified, and *FLAD1* expression is upregulated in various tumor types, especially in hypoxic tumors. *FLAD1* depletion compromises tumor’s adaptation to hypoxia by disrupting mitochondrial complex II activity, leading to an imbalance between succinate and fumarate, and consequent failure to adapt to hypoxia. Subsequently, we identified a drug-like inhibitor of FLAD1, which selectively inhibits the growth of tumor cells under hypoxia. Our findings reveal FLAD1 as an innovative therapeutic target for hypoxic tumors.

## Introduction

Hypoxia is a hallmark of the tumor microenvironment, with tumor survival under hypoxic conditions reflecting the remarkable adaptabilities of cancer cells(Keith & Simon, 2007; Chen *et al*, 2023). Hypoxia presents a significant metabolic challenge to tumors; paradoxically, however, tumors not only survive but often thrive under these conditions. The success of this adaptability is partly due to the metabolic flexibility and plasticity of cancer cells, which confers a survival advantage under stress.

Advancements in hypoxia research have enhanced our understanding of the tumor microenvironment, particularly the contribution of the HIF pathway to the hypoxic adaptation of cancer cells(Semenza *et al*, 1997; Maxwell *et al*, 1999; Wang *et al*, 1995; Iliopoulos *et al*, 1996). Despite the potential of targeting hypoxic tumors in cancer therapy through strategies such as hypoxia-activated prodrugs, HIF pathway inhibitors, and anti-angiogenic agents, the therapeutic outcomes remain modest(Badar *et al*, 2016; Tap *et al*, 2017; Bui *et al*, 2022; Bellou *et al*, 2013; Ebos & Kerbel, 2011; Shojaei, 2012; Chen & Cleck, 2009). While the prospect of targeted therapy for hypoxic tumors is promising, it currently lacks established clinical practice. This limited efficacy is attributable to several factors, one of which is the insufficient understanding of the mechanisms underlying tumor adaptation to hypoxia, which hampers the development of safe and effective targeted therapies.

Under hypoxic conditions, tumor cells undergo metabolic reprogramming to reshape their energy metabolism pathways for adaptation. However, this also causes them to be dependent on specific metabolic pathways. This phenomenon offers potential opportunities for identifying metabolic vulnerability targets in tumors. Nevertheless, the complexity of cellular metabolism and the multifaceted interactions between cancer cells and the tumor microenvironment (TME) present significant challenges in identifying metabolic targets in hypoxic tumors(Elia & Haigis, 2021; Xiao *et al*, 2023). Traditional experimental approaches such as CRISPR screening are beset by several limitations: they are time-consuming and labor-intensive; the data generated often pertain to a specific tumor cell line, yet tumor metabolism is characterized by heterogeneity, rendering such results potentially biased; moreover, the hypoxic microenvironments simulated *in vitro* may not accurately reflect the actual TME(Liu *et al*, 2023; Kang *et al*, 2020). To address these issues, we leveraged the transformer-based model, Geneformer, pre-trained on about 30Lmillion single-cell transcriptomes to learn meaningful representations, which offer a profound understanding of gene networks(Theodoris *et al*, 2023). Pre-trained foundation models enable unparalleled insight into the intricate dynamics of genetic interactions, facilitating precise predictions in the complex landscape of network biology. However, while Geneformer excels in network dynamics understanding, it is not directly applicable for predicting metabolic vulnerabilities in hypoxic tumors. Building upon Geneformer’s architecture, we have fine-tuned a specialized deep learning model, termed DepFormer. DepFormer enables the utilization of real tumor tissue data sourced from clinical patients, facilitating the prediction of metabolic targets that tumors depend upon under hypoxic conditions with enhanced precision and relevance.

Herein, we reveal a hitherto unexplored link between *FLAD1* and the hypoxic adaptation of cancer cells through comprehensive metabolic network analysis, deep learning model construction, *in silico* perturbation analysis and rigorous experimental validation. FLAD1 catalyzes the formation of flavin adenine dinucleotide (FAD), and loss of function mutations in *FLAD1* lead to a human genetic disease named multiple acyl-CoA dehydrogenase deficiency (MADD), which is characterized by the FAD-related defects in mitochondrial metabolism(Olsen *et al*, 2016; Ryder *et al*, 2018). Germline mutations in the alpha and beta subunits of electron transfer flavoprotein (*ETFA* and *ETFB*), and electron transfer flavoprotein dehydrogenase (*ETFDH*) can also lead to MADD, and FAD is the coenzyme for ETFA, ETFB and ETFDH (Mereis *et al*, 2021). However, the roles of *FLAD1* and FAD in hypoxic tumors remain unknown. By examining the roles of FLAD1 and its product FAD in the electron transport chain under hypoxia, especially their critical role in maintaining cellular metabolism under low oxygen conditions, we provide new insights into the metabolic adaptation of cancer and the potential targeting of hypoxic tumors.

## Results

### Systematic analysis of the metabolic reprogramming in hypoxic tumor cells

Hypoxia is a hallmark of solid tumors and poses a significant challenge for cancer therapy. Here we aim to identify the metabolic genes critical for tumor hypoxic adaptation, and targeting these metabolic genes could selectively inhibit the proliferation or survival.

We investigate the metabolic reprogramming of tumor cells by comparing the metabolic enzyme network difference between hypoxic and non-hypoxic samples. The cancer genome atlas (TCGA) sample specific genome-scale metabolic models (GEMs) were retrieved from Francesco Gatto et al work(Gatto *et al*, 2020), which contains 4849 TCGA tumor samples, and then converted GEMs to metabolic enzyme networks. The hypoxic status of each sample is defined by Buffa hypoxia score(Buffa *et al*, 2010).

We first calculated the similarity between metabolic network embedding obtained by the graph2vec algorithm for different samples(Narayanan *et al*, 2017). In lung cancer, we found that the similarity of metabolic network among samples within the same hypoxic state was significantly higher than that in samples in different hypoxic states, including hypoxic and non-hypoxic states (**Extended Data Fig. 1a**). We also investigated genes based on their network centrality, given that central nodes tend to act as hubs with higher biological importance. The centrality metrics include degree centrality, betweenness centrality, eigenvector centrality and closeness centrality (**Methods**). We generated gene-wise vectors of centrality measures for each sample specific enzyme network and calculated the Euclidean distance between these vectors. Similarly, we found that in lung cancer, the distance between metabolic networks of samples with the same hypoxic state was significantly lower than that between samples with different hypoxic states (**Extended Data Fig. 1b-d**). These results suggest that lung cancer shows significant hypoxia associated metabolic reprogramming.

Furthermore, for each gene, we compared the differences in the centrality metrics between hypoxic and non-hypoxic tumors, and identified genes with at least one centrality metrics significantly up-regulated in hypoxic tumors (Wilcoxon test adjusted P value < 0.05). Then these genes were selected to perform metabolic pathway enrichment analysis. We found that the oxidative phosphorylation (OXPHOS) pathway was the most significantly enriched pathway (**Fig. 1a**), suggesting that OXPHOS related genes may have important functions in the metabolic adaptation under hypoxia.

**Fig. 1.**
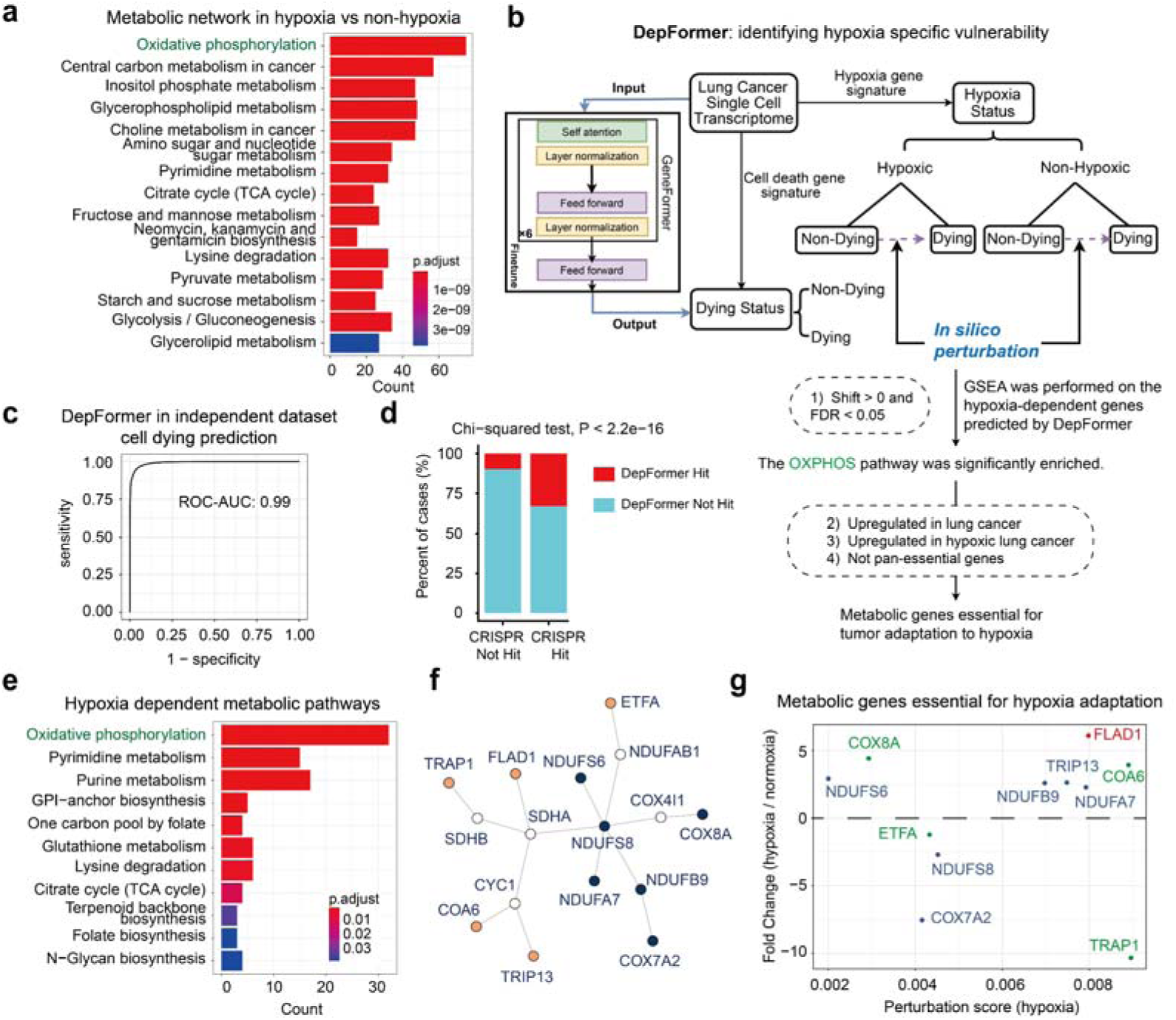
Metabolic dependencies of hypoxic tumor cells revealed by network analysis and deep learning. **a,** The results of metabolic pathway over-representation enrichment analysis using the gene nodes with up-regulated centrality in the enzyme network of hypoxic tumor compared with non-hypoxic tumor. Genes with at least one centrality metrics significantly up-regulated in hypoxic tumors (Wilcoxon test adjust P value < 0.05) were included for analysis. Centrality metrics of gene nodes in enzyme network include degree centrality, betweenness centrality, eigenvector centrality and closeness centrality. **b**, Workflow of *in silico* perturbation using a fine-tuned Geneformer, named DepFormer, to identify metabolic dependencies of hypoxic tumor cells. We first used the activity of cell death-related and hypoxia-related gene sets to divide cells into dying cells/non-dying and hypoxic/non-hypoxic cells based on single-cell data, and then fine-tune the foundational model GeneFormer to predict cell dying state. We then identified the genes whose *in silico* deletion in hypoxic cells or non-hypoxic cells significantly shifted the DepFomer cell embeddings from non-dying cell state towards the dying cell state. By comparing the differences of *in silico* perturbation results between hypoxic and non-hypoxic cells, we can identify metabolic genes that are specifically dependent in hypoxic tumors. **c**, The performance of fine-tuned model shown as receiver operating characteristic curve (ROC) and corresponding area under curve (AUC) score on the independent test dataset from Qian, J. et al. **d**, Validation of *in silico* perturbation results using cell line CRISPR screening. The CRISPR-Cas9 screen data of 81 lung cancer cell lines was downloaded from Depmap project (23Q4 version). The dependency probabilities value is between 0 and 1. Genes with a dependency score > 0.8 were marked as cell dependent, genes that were dependent in more than 20 cell lines were marked as CRISPR hit, and other genes were marked as CRISPR not-hit. *In silico* perturbation hit genes were defined as the effect value was > 0, and the FDR was < 0.05. **e**, The results of metabolic pathway over-representation enrichment analysis using genes predicted to be specifically required for the survival of hypoxic tumor cells. **f**, The minimal subgraph of the protein-protein interaction network of oxidative phosphorylation pathway related genes. This minimal subgraph contains 11 genes (colored in orange and blue), complete oxidative phosphorylation pathway related gene protein-protein interactions network is shown in **Extended Data Fig. 4a**. We extracted the smallest subgraph connecting these 11 gene nodes by minimum spanning tree based approximation (KB) algorithm. The orange dots represent genes that are not involved in the oxidative phosphorylation pathway directly but interact with oxidative phosphorylation-related genes. The deep blue dots indicate genes that are part of the oxidative phosphorylation pathway. **g**, DepFormer identified OXPHOS-related genes involved in tumor hypoxia adaptation. The perturbation score reflects the importance of the predicted genes. The fold change indicates the selectivity of gene perturbations in hypoxic tumors.

### Deep learning reveals the metabolic dependencies of hypoxic tumor cells

Among all pan-cancer types, lung cancer shows the most significant difference in the metabolic enzyme networks between hypoxic and non-hypoxic samples. Here, we fine-tuned a deep learning model, “DepFormer”, to predict the hypoxia-dependent metabolic genes using lung cancer single cell RNA-seq data (**Fig. 1b**). We first defined a “cell dying” status based on the expression of 12 cell death-related gene sets(Zou *et al*, 2022). The definition of the “cell dying” status enables the DepFormer model to identify genes that are critical for cell survival under hypoxic conditions following *in silico* perturbations. AUCell (Aibar *et al*, 2017) was used to calculate the activity score of these gene set for each tumor cell, referred to as the “cell dying score”. In the dataset obtained from Prazanowska et al (Prazanowska & Lim, 2023), the cell dying scores were quantified for 42529 tumor cells, which showed an apparent bimodal distribution (**Extended Data Fig. 2a**). Thus, the Gaussian mixture model (GMM) is applied to classify these cells into two types: dying cell and non-dying cell. We then fine-tuned the GeneFormer model (Theodoris *et al*, 2023), a foundational transformer model pre-trained on a large-scale corpus of approximately 30 million single cell transcriptome to enable context-aware predictions in settings with limited data in network biology, to learn the dying and non-dying status of cells. The fine-tuned Geneformer, named DepFormer, can accurately classify each single cell as either dying or non-dying status with high accuracy (AUC=0.99) in an independent lung cancer single cell dataset(Qian *et al*, 2020) (**Fig. 1c** and **Extended Data Fig. 2b**). We then identified the genes whose *in silico* deletion significantly shifted the DepFormer cell embeddings from the non-dying cell state towards the dying cell state. These results indicate that cell survival depends on specific genes. To validate this approach in identifying critical cell dependency genes, we compared the results with CRISPR screening in lung cancer cells from DepMap dataset(Tsherniak *et al*, 2017). Indeed, genes identified by DepFormer *in silico* perturbation are significantly enriched in CRISPR hits (**Fig. 1d** and **Extended Data Fig. 3**).

To identify genes specifically required under hypoxic conditions, we used Gaussian mixture model to classify the cells into hypoxic and non-hypoxic states based on the score of hypoxia-related gene sets quantified by AUCell. Subsequently, *in silico* perturbation was conducted separately in these two cell states. We further conducted gene enrichment analysis on the output of DepFormer. The metabolic genes required for hypoxia adaptation, as predicted by DepFormer, are significantly enriched in the OXPHOS pathway (**Fig. 1e**), which is consistent with the findings presented in **Figure 1a**. This suggests that the OXPHOS pathway may play a crucial role in the hypoxia adaptation of tumors. We further filtered the OXPHOS-related genes (**Extended Data Fig. 4a**) based on the following four criteria to identify metabolic genes that are crucial for tumor hypoxic adaptation: 1) genes that are predicted by DepFormer as hypoxic lung cancer cell dependent; 2) genes that are significantly up-regulated in lung cancer compared to normal lung tissues; 3) genes that are significantly up-regulated in hypoxic lung cancer relative to non-hypoxic lung cancer; 4) genes that are not classified as pan-essential (i.e., essential for all cell types) (**Fig. 1b**). Ultimately, we identified 11 candidate metabolic genes that are essential for tumor adaptation to hypoxia (**Extended Data Fig. 4b and Fig. 1f**).

Notably, *FLAD1* is not categorized as a pan-essential gene. It exhibits a high perturbation score in hypoxic lung cancer cells, but not in non-hypoxic lung cancer cells (**Fig. 1g and Extended Data Fig. 4c**). In these identified 11 genes, *TRAP1*, *COA6*, *ETFA*, and *COX8A* have been previously reported to be associated with hypoxic adaptation and the HIF pathway (Bruno *et al*, 2022; Shekhova *et al*, 2019; Yan *et al*, 2022; Chen *et al*, 2020), suggesting the robustness of this analysis in identifying hypoxia adaptation related genes. However, *FLAD1* has not been characterized in this context previously. As a leading candidate among hypoxia-dependent gene cohorts, FLAD1 represents a promising therapeutic target for hypoxic tumors.

Our research further demonstrates that both mRNA and protein expression levels of *FLAD1* are significant upregulated in tumors compared with normal control tissues (**Fig. 2a, b** and **Extended Data Fig. 5a**). This upregulation of FLAD1 expression is also evident in hypoxia tumors compared with non-hypoxic tumors (**Fig. 2c**). Independent dataset within the Oncomine database further corroborate a pronounced elevation of *FLAD1* in lung cancer compared with normal lung tissue (**Extended Data Fig. 5b**). Among patients with lung adenocarcinoma (LUAD), a substantial increase in the copy number of *FLAD1* is evident, with approximately 20% of patients exhibiting amplification of *FLAD1* DNA. This percentage increases to 26% in hypoxic tumors, surpassing the 11.7% observed in non-hypoxic tumors. At the pan-cancer level, we also observed a significant amplification in DNA copy number of the *FLAD1* gene across tumor samples, particularly pronounced in hypoxic tumors (**Fig. 2d**). Moreover, analyses of data from four independent studies indicate that the abundance of FAD is also increased in tumors(Goldberg *et al*, 2021; Moreno *et al*, 2018; Jung *et al*, 2013) (**Fig. 2e**). In summary, our findings highlight a significant upregulation of *FLAD1* levels in tumors, particularly in hypoxic tumors, suggesting its potential involvement in the adaptive response to tumor hypoxia. Additionally, in the TCGA pan-cancer dataset and specific tumor types such as KIRC, LGG, LIHC, and SKCM, high expression levels of *FLAD1* portends adverse overall survival rates (**Fig. 2f** and **Extended Data Fig. 5c**), highlighting its prognostic significance.

**Fig. 2.**
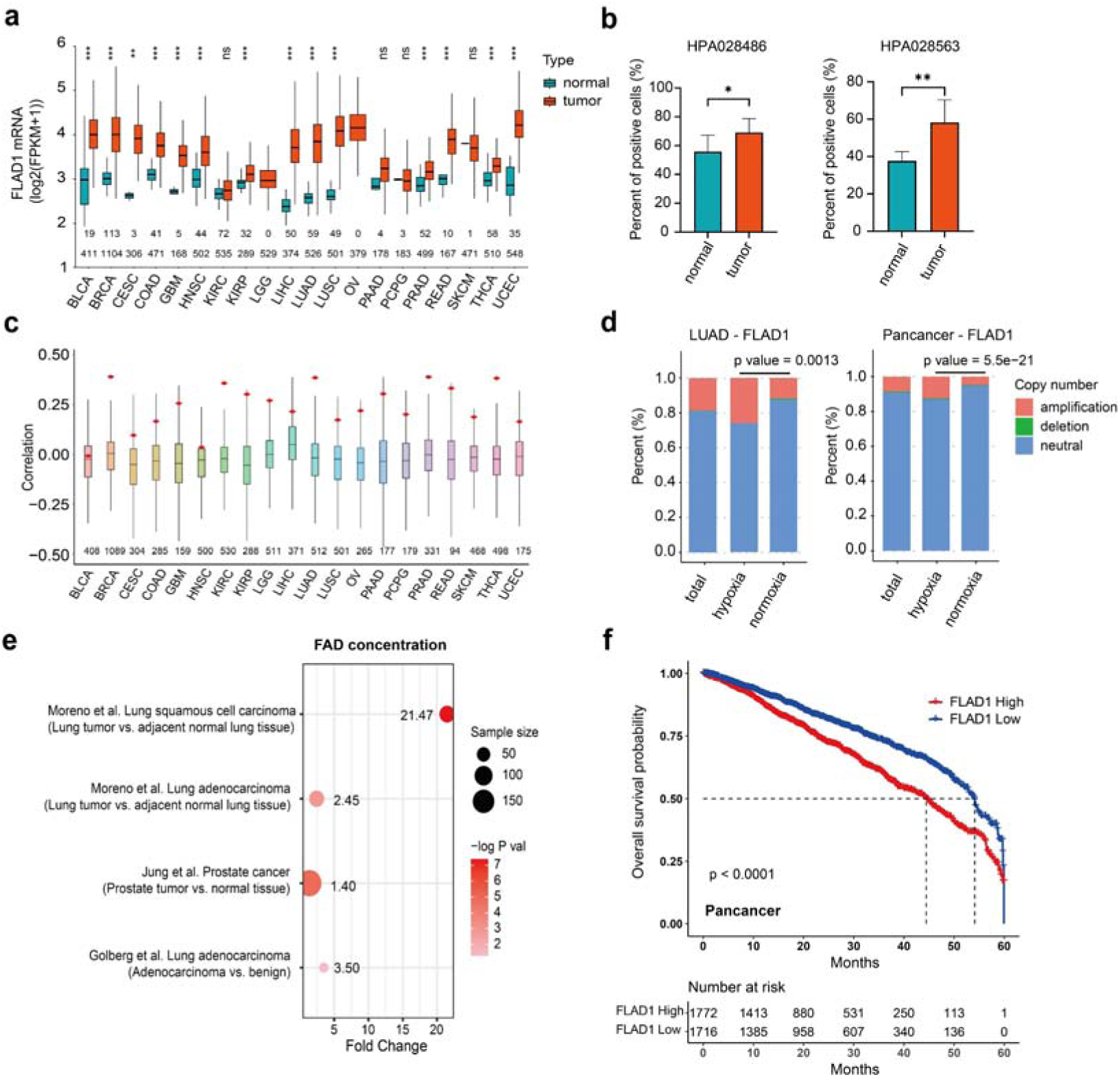
Characteristics of FLAD1 in human tumors. **a**, Comparison of FLAD1 mRNA levels between various types of tumors and corresponding normal tissues in the TCGA database. The numbers above the x-axis indicate the count of normal tissue samples (first row) and tumor samples (second row). Data are presented as median (Q1, Q3). **b**, Immunohistochemical results for FLAD1 protein levels quantified in lung cancer from the HPA database, using two different FLAD1-targeting antibodies, HPA028486 and HPA028563. **c**, Correlation between gene mRNA expression levels and Buffa hypoxia scores across various tumor types in the TCGA database. The red dots signify the correlation of *FLAD1* mRNA levels with Buffa hypoxia scores. The numbers above the x-axis denote the count of tumor samples analyzed. Data are depicted as median (Q1, Q3). **d**, Variations in *FLAD1* DNA copy number among normal lung tissue, hypoxic lung cancer, and non-hypoxic lung cancer samples (left panel), as well as in pan-cancer level including tumor, hypoxic tumor, and non-hypoxic tumor samples (right panel), in the TCGA database. The statistical significance of differences in *FLAD1* copy number amplification between hypoxic and non-hypoxic tumor samples was assessed using Fisher’s exact test. **e**, Content of FAD in tumors across four independent datasets. **f**, Kaplan-Meier curves of overall survival for pan cancer with high and low *FLAD1* expression in TCGA database. The “Number at risk” represent the total number of survivors at the beginning of each specified period (in months). The top row represents the number of patients with high *FLAD1* expression, and the bottom row represents the number of patients with low *FLAD1* expression.

### FLAD1 is required for tumor hypoxia adaptation

To verify the functional role of FLAD1 in hypoxic adaptation, the CRISPR-Cas9 system was used to knockout the FAD synthesis domain of FLAD1 (**Extended Data Fig. 6a**). A rigorous screening and evaluation process was then performed, including both PCR and Western blot, to confirm the successful depletion of *FLAD1* in two tumor cell lines, PC9 and H520 (**Extended Data Fig. 6b, c**). Knockout of the *FLAD1* gene resulted in a significant decrease in the intracellular FAD levels (**Extended Data Fig. 6d**). Cell proliferation assays revealed that, compared to the wild-type counterparts, *FLAD1*-null H520 and PC9 lung cancer cells exhibited a notable suppression of proliferation under hypoxic conditions (1% O2) (**Fig. 3a**).

**Fig. 3.**
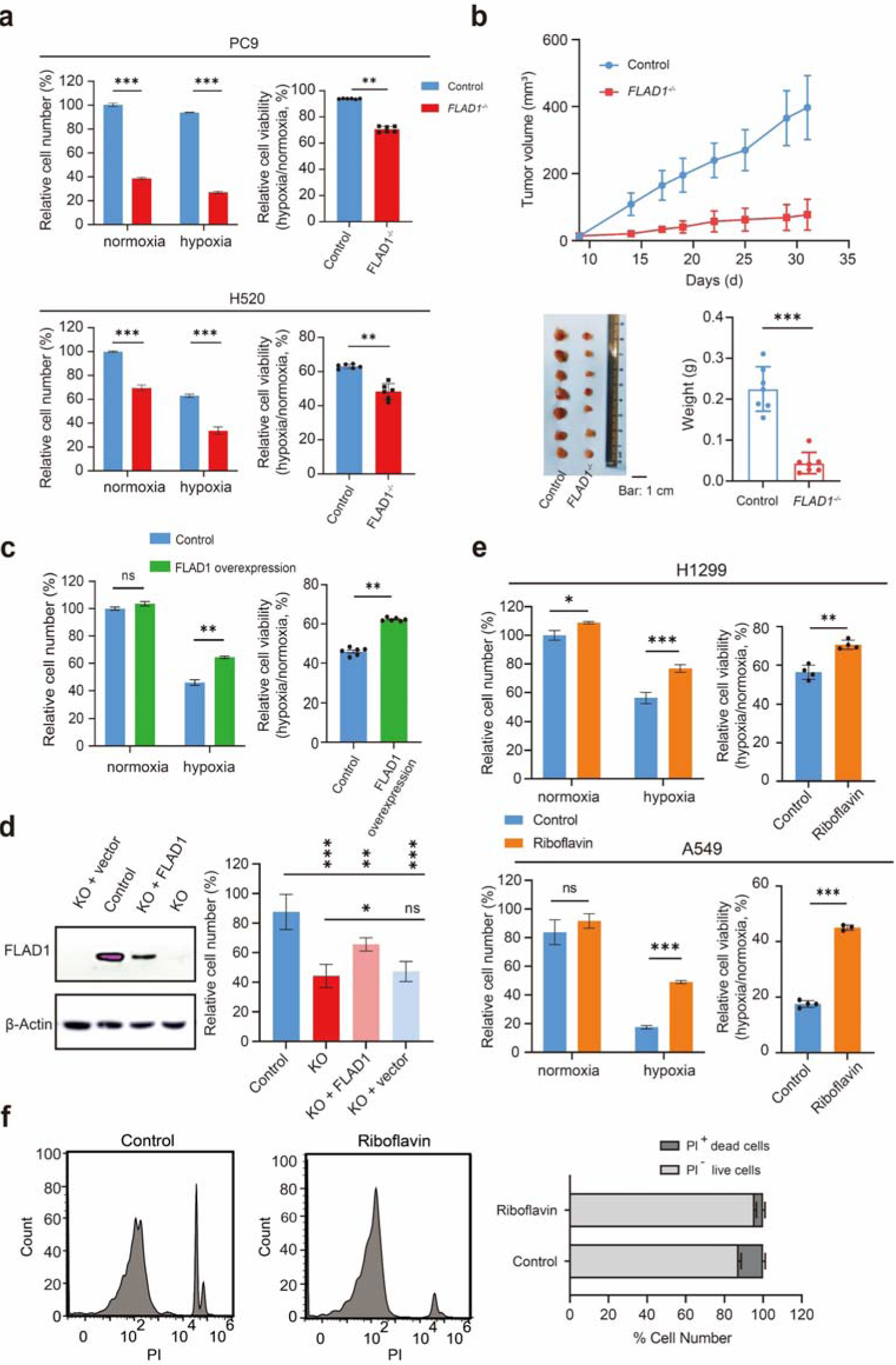
FLAD1 knockout impairs, while its overexpression enhances cancer cell adaptation to hypoxia. **a**, Proliferation assays of PC9 (top) and H520 (bottom) tumor cells, comparing control groups with *FLAD1* knockout cells under normoxic (21% O2) and hypoxic (1% O2) conditions, where cell viability (%) = 100% × (relative quantity of cells under hypoxia / relative quantity of cells under normoxia). **b**, Tumor volume at 35 days post-injection of wild-type and *FLAD1* knockout PC9 cells subcutaneously into nude mice is shown (top), along with a comparison of tumor size and weight (bottom); scale bar: 1 cm. **c**, Proliferation assays of control and *FLAD1*-overexpressing A549 cells under normoxic and hypoxic (1% O2) conditions, where cell viability (%) = 100% × (relative quantity of cells under hypoxia / relative quantity of cells under normoxia). **d**, Lentivirus-mediated expression of *FLAD1* can partially restore the proliferative capacity of PC9 tumor cells with *FLAD1* knockout. KO: *FLAD1* knockout; FLAD1: lentivirus-mediated expression of *FLAD1*; Vector: lentivirus-mediated negative control. **e**, Bar graph indicating the proliferation of H1299 and A549 cells after supplementation with 0.4 uM riboflavin under normoxic (21% O2) and hypoxic (1% O2) conditions. **f**, PI staining demonstrating cell death in control and 0.4 uM riboflavin-supplemented H1299 cells. PI: Propidium Iodide.

While previous study employing CRISPR screening have identified several genes, including *FLAD1*, that may play roles in the hypoxic adaptation of tumor cells(Garcia-Bermudez *et al*, 2022), the function of the *FLAD1* gene was not highlighted. Our study unveils, for the first time, the pivotal role of the *FLAD1* gene in the adaptation of tumor cells to hypoxic conditions. Gene knockout experiments demonstrate that tumor cells lacking *FLAD1* exhibit marked intolerance to low oxygen levels, underscoring its unique and previously unrecognized function in hypoxic adaptation. Furthermore, a xenograft experiment was conducted by subcutaneously inoculating both *FLAD1*-knockout and wild-type PC9 cells into nude mice. After a 35-days observation period, we performed a comparative analysis of tumor volume and weight in the murine hosts. Intriguingly, the tumorigenic capacity of the *FLAD1*-knockout cells was significantly diminished *in vivo*, as depicted in **Fig. 3b**.

Subsequently, lentivirus-mediated *FLAD1*-overexpressed A549 tumor cells were constructed **(Extended Data Fig. 6e)**. Remarkably, *FLAD1* overexpression elevated the proliferation rate of A549 cells under hypoxic conditions (**Fig. 3c**). Re-expressing *FLAD1* in *FLAD1*-knockout PC9 tumor cells partially rescued their proliferative defects (**Fig. 3d**). The *FLAD1* gene plays a crucial role in the riboflavin metabolism. Within cells, riboflavin is first converted to flavin mononucleotide (FMN), and then FLAD1 catalyzes the conversion of FMN to FAD (Torchetti *et al*, 2010; Barile *et al*, 2016). Supplementation of H1299 and A549 cells with riboflavin mitigated the deleterious effects of hypoxia on these tumor cells. This protective effect is manifested in several specific ways. Notably, under hypoxic conditions, cells exhibited better-preserved morphology, enhanced proliferation rates, and reduced apoptotic levels (**Fig. 3e, f and Extended Data Fig. 6f**). Collectively, the outcomes from both *in vitro* and *in vivo* experiments corroborate and reinforce the assertion that FLAD1 is intricately involved in the adaptive response of tumor cells to hypoxic conditions.

Given that FLAD1 is implicated in riboflavin metabolism, we embarked on a specialized investigation to ascertain whether this metabolic pathway also holds significance in the context of tumor hypoxia adaptation. Leveraging information from the KEGG and Reactome database, we depicted the riboflavin metabolic pathway. As illustrated in **Extended Data Fig. 7a,** genes *SLC52A1, SLC52A2, SLC52A3, RFK* and *FLAD1* are involved in the synthetic metabolism of FAD. Through analysis of gene expression data from tumor patients in the TCGA database, we conducted comprehensive comparisons of tumor and normal tissues, as well as hypoxic and non-hypoxic tumor subgroups. Our analysis unveiled notable trends within the FAD metabolic pathway-related genes. Specifically, genes associated with FAD synthetic metabolism exhibited elevated mRNA levels in tumors, particularly in the hypoxic tumor subset (**Extended Data Fig. 7a, b**). Tumors were stratified into five distinct hypoxic subtypes— “Very Low”, “Low”, “Moderate”, “High” and “Very High”—based on their hypoxia scores. It was found that an increase in hypoxia levels correlated with elevated riboflavin metabolism, as evidenced in **Extended Data Fig. 7c**. This implies a metabolic shift favoring FAD production in hypoxic tumors. Lumiflavin, a riboflavin analog, impedes the cellular uptake of riboflavin, disrupting riboflavin metabolism. Cells treated with lumiflavin exhibited selective inhibition under hypoxic conditions (**Extended Data Fig. 7d**). Moreover, restoration of FAD partially rescued proliferation of *FLAD1*-knockout PC9 cells (**Fig. 3d**). These findings underscore the importance of FAD and riboflavin metabolism in tumor hypoxic adaptation.

### Mitochondrial metabolic reprogramming in FLAD1-mediated tumor hypoxia adaptation

Loss-of-function mutations in *FLAD1* in humans lead to MADD genetic disease, characterized by symptoms related to mitochondrial metabolism defects, such as dysphagia, muscular weakness, and respiratory distress (**Extended Data Fig. 8a**)(van Rijt *et al*, 2019; Brandão *et al*, 2021; Béhin *et al*, 2016). We hypothesize that FLAD1 facilitates tumor adaptation to hypoxic conditions by reprogramming mitochondrial metabolism. We categorized TCGA tumors into two groups based on *FLAD1* expression levels: high and low *FLAD1* expression. Using Gene Ontology (GO) analysis to compare genes upregulated in the high *FLAD1* expression group to those in the low expression group, we found a significant association with mitochondrial components. Notably, four of the top ten enriched cellular components pertain to mitochondria (**Extended Data Fig. 8b**).

Immunofluorescence localization further confirmed the co-localization of FLAD1 with mitochondria (**Extended Data Fig. 8c**). Comparing reactive oxygen species (ROS) levels between wild-type and *FLAD1*-knockout cells, we observed elevated ROS levels to varying degrees in *FLAD1*-knockout A549 and PC9 cells (**Extended Data Fig. 8d**). ROS are primarily produced by mitochondria, suggesting that the absence of *FLAD1* leads to mitochondrial dysfunction and subsequent ROS overproduction (Kowalczyk *et al*, 2021; Choi *et al*, 2024; Palma *et al*, 2024). Subsequent experiments were conducted to investigate the impact of *FLAD1* knockout on mitochondrial function. We examined the effects of *FLAD1* knockout on cellular respiratory function and mitochondrial membrane potential. Knockout of *FLAD1* in PC9 and H520 cells resulted in decreased basal respiration levels and spare respiratory capacity (SRC), along with reduced mitochondrial membrane potential, indicating impaired mitochondrial function (**Fig. 4a, b**).

**Fig. 4.**
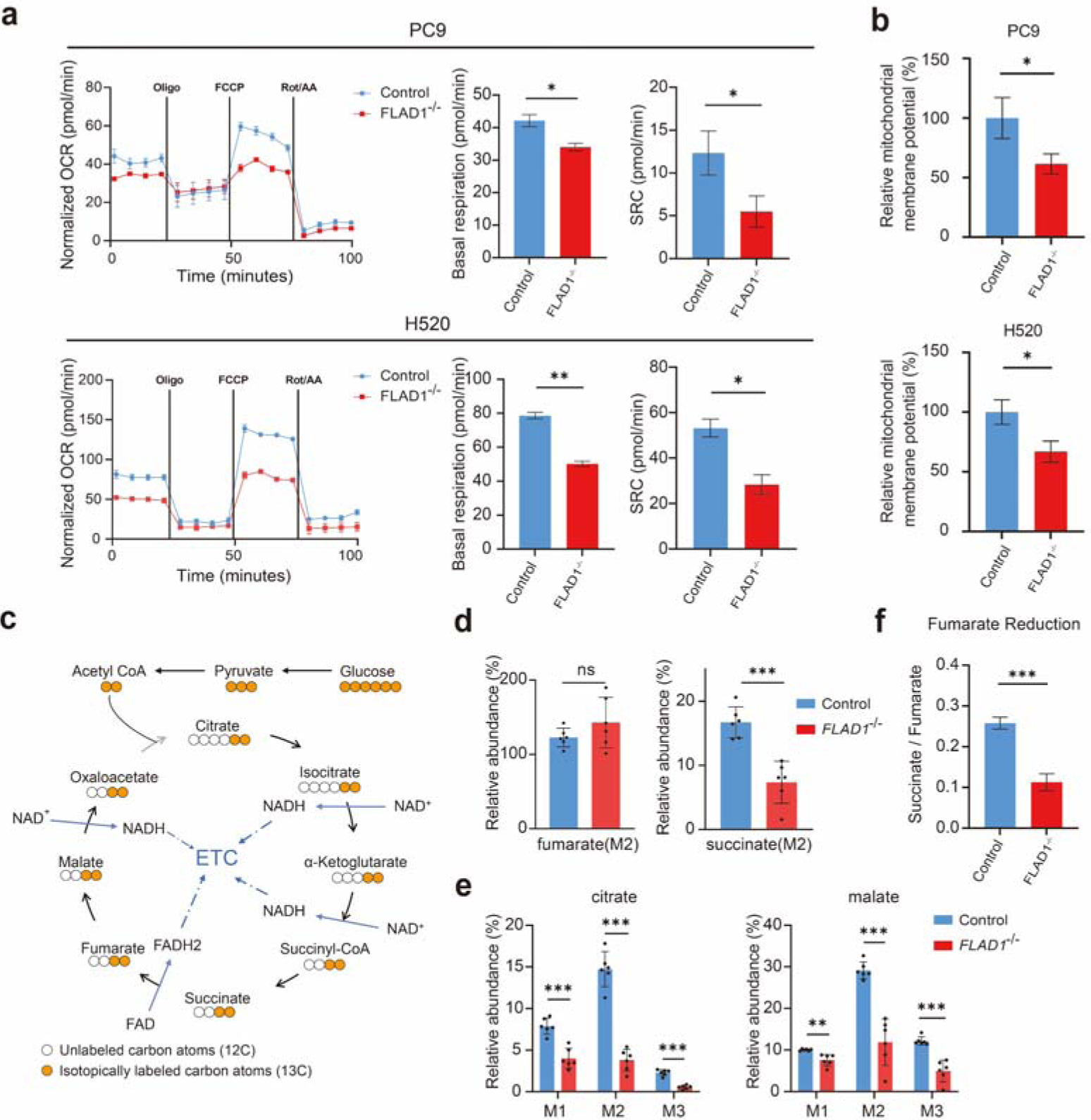
Loss of *FLAD1* impairs mitochondrial metabolic activity. **a**, Mitochondrial seahorse assays of tumor cell lines PC9 and H520. The oxygen consumption rate (OCR) is measured for control and *FLAD1* knockout (KO) cells under normoxia, with quantification of basal respiration and spare respiratory capacity (SRC) in both control and *FLAD1* knockout groups. **b**, Bar graphs illustrate the variance in mitochondrial membrane potential between H520 and PC9 cells under normoxia following *FLAD1* knockout compared to their respective control groups. **c**, Intermediates of the mitochondrial TCA cycle in PC9 tumor cells labeled with the isotope 13C-glucose. Hollow circles represent unlabeled carbon atoms (12C), while orange circles represent isotope-labeled carbon atoms (13C). **d**, Relative levels of 13C glucose-labeled fumarate and succinate in control and *FLAD1* knockout PC9 cells under hypoxic conditions (1% O2). **e**, Relative levels of 13C-glucose-labeled TCA cycle intermediates in control and *FLAD1* knockout PC9 cells under hypoxic conditions. M, unlabeled mass of isotope; Mn, native metabolite mass (M) plus number of isotopically labeled carbons (n). **f**, The reduction level of fumarate in wild-type and *FLAD1* knockout PC9 cells under hypoxia. The value is represented by the ratio of succinate (M+3) to fumarate (M+3).

Mitochondria play two principal roles within the cell: first, as the cell’s “powerhouse,” they are involved in the synthesis of ATP, the cell’s energy currency; second, they are implicated in the synthesis of various metabolic substances, with the tricarboxylic acid (TCA) cycle being a critical metabolic pathway. The TCA cycle is central to cellular metabolism; it provides substantial energy through the oxidation of fuel molecules and contributes to synthesizing precursors for numerous biomolecules. Utilizing targeted metabolomics with labeled 13C-glucose, we observed a decrease in the levels of TCA cycle intermediates in *FLAD1*-knockout cells under hypoxia (**Fig. 4c, d, e**). This implies that the suppression of growth in *FLAD1*-knockout cells under hypoxia is likely due to a blockade in mitochondrial metabolism, specifically in the TCA cycle. The TCA cycle in mitochondria functions akin to a mechanical gear, engaging with various metabolic pathways within mitochondria, much like gears intermeshing with one another. This raises the question: why does the metabolic “gear” of *FLAD1*-knockout cells get stuck under hypoxic conditions?

The product of FLAD1, FAD, is a crucial cofactor that serves as an intermediate for electron transfer across various proteins. The four high-risk genes associated with the genetic disorder MADD are *FLAD1, ETFA, ETFB* and *ETFDH*. These genes are closely associated with intracellular FAD homeostasis. Therefore, we hypothesize that FLAD1 primarily participates in tumor hypoxia adaptation through its product, FAD. In our deep learning model, we discerned that hypoxic cells rely on OXPHOS, which consists of two parts: the electron transport chain (ETC) and chemiosmosis. Complex II of the ETC is a prototypical FAD-dependent flavoprotein. Previous reports indicate that under hypoxic conditions, the activity of mammalian complex II can reverse, with fumarate serving as the terminal electron acceptor in the ETC, being reduced to succinate by complex II(Kita *et al*, 2002; Lemire & Oyedotun, 2002; Maklashina *et al*, 1998; Spinelli *et al*, 2021) (**Extended Data Fig. 9a**). This specialized electron transport process occurs exclusively under low oxygen conditions and is coupled to the TCA cycle. Hence, we hypothesize that the “gear” that becomes stuck in *FLAD1*-knockout cells may be located at complex II. Significantly, within our deep learning model, multiple subunits of complex II also demonstrated a dependency for cellular survival under hypoxic conditions (**Extended Data Fig. 9b**).

Treatment of both *FLAD1* wild-type and knockout cells with complex II inhibitors (Thenoyltrifluoroacetone, TTFA and 3-nitropropionic acid, 3-NPA) revealed that *FLAD1*-knockout cells no longer exhibited sensitivity to hypoxia compared to wild-type cells upon inhibition of complex II (**Fig. 5a**). Conversely, cells with inhibited complex II demonstrated sensitivity to hypoxia (**Fig. 5a**). qPCR validation indicated that *FLAD1* knockout did not affect the expression of the four principal subunits of complex II (**Extended Data Fig. 9c**); however, *FLAD1* knockout resulted in a decline in complex II function (**Fig. 5b**). The metabolomic analysis further supports the observation of altered complex II activity in *FLAD1*-knockout cells, manifesting as a decrease in the reverse activity, specifically the reduction of fumarate to succinate (**Fig. 4f**). Comprehensive metabolic profiling was conducted through untargeted metabolomics to examine the overall metabolic landscape of cells (**Extended Data Fig. 9d**). Clustering results indicated distinct metabolic pathway alterations between the *FLAD1* knockout and control groups (**Extended Data Fig. 9e**). Enrichment analysis of the downregulated metabolites in the *FLAD1* knockout group revealed significant downregulation across multiple metabolic pathways (**Fig. 5c**). From these observations, we conclude that *FLAD1* knockout leads to a blockade of the mitochondrial TCA cycle under hypoxic conditions, which is associated with reduced activity of complex II. Complex II relies on FAD to transfer electrons from the ETC to fumarate. This transfer not only effectively neutralizes harmful electrons but also facilitates the recycling of substrates back into the TCA cycle, thereby propelling the metabolic “gear” forward (**Fig. 5d**).

**Fig. 5.**
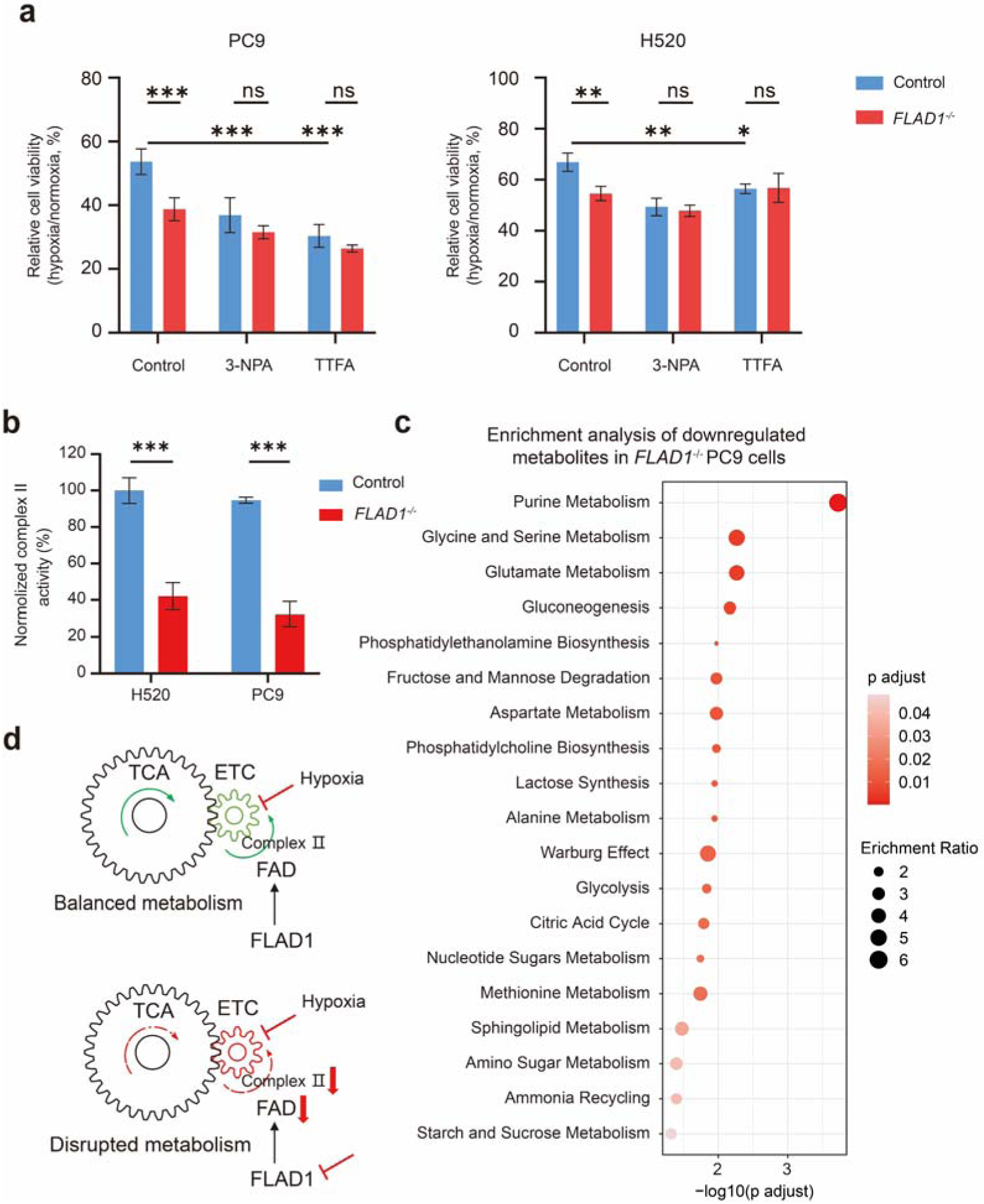
Mitochondrial complex Ⅱ and FLAD1 mediated hypoxia adaptation. **a,** The impact of complex II inhibitor (3-NPA, TTFA) on the proliferation of control and *FLAD1* knockout PC9 and H520 cells. By comparing the relative cell number under hypoxic conditions to that under normoxic conditions and normalizing the data, the percentage of viable cells can be quantified. For PC9 cells, treat with 15 uM 3-NPA and 200 uM TTFA. For H520 cells, treat with 30 uM 3-NPA and 150 uM TTFA. **b**, Comparison of complex II activity in PC9 and H520 cells between wild-type and *FLAD1* knockout groups under normoxia. **c**, Enrichment results of significantly down-regulated metabolites in pathways within the *FLAD1* knockout group under hypoxia. Exploring the metabolites of wild type and knockout groups through non-targeted metabolomics, selecting significantly down-regulated genes in the knockout group, and conducting pathway enrichment analysis using MetaboAnalyst algorithm. **d**, Mechanism for mitochondria complex II and FLAD1 mediated tumor hypoxia adaptation. Under hypoxia, the electron transport chain lacks sufficient O2 as the final electron acceptor. At this time, mitochondrial metabolism requires an adequate supply of FAD to maintain the reverse process of mitochondrial complex II activity, which involves the conversion from succinate dehydrogenase activity to fumarate reductase activity, relying on FAD as a cofactor. Deletion of FLAD1 results in impairment of complex II activity in hypoxic environments, which in turn interferes with multiple metabolic pathways coupled to the electron respiratory chain (such as the TCA cycle).

To investigate whether similar patterns exist in human cancer patients, we stratified tumors from the TCGA dataset into high and low *FLAD1* expression groups. Gene set enrichment analysis (GSEA) indicated that in the subset with high *FLAD1* expression, there is an upregulation of multiple metabolic pathways, particularly pronounced for pyrimidine metabolism and OXPHOS (**Extended Data Fig. 10a**). In de novo pyrimidine biosynthesis, dihydroorotate dehydrogenase (DHODH) catalyzes the oxidation of dihydroorotate to orotate, concomitantly transferring electrons to the ETC. This process is therefore highly coupled to the ETC. Furthermore, analysis across various tumors types in the TCGA dataset revealed that genes related to OXPHOS were expressed at significantly higher levels in tumors with high *FLAD1* expression (**Extended Data Fig. 10b**). GSEA demonstrated that genes associated with mitochondrial OXPHOS were highly active in lung cancer patients with elevated *FLAD1* expression levels (**Extended Data Fig. 10c**). These findings suggest a close relationship between FLAD1 and mitochondrial function, highlighting FLAD1’s role in influencing mitochondrial complex Ⅱ activity to facilitate tumor adaptation to hypoxia. The characterized metabolic profiles and the genetic analyses support a model in which FLAD1 influences overall mitochondrial efficiency and the metabolic adaptation of tumors to oxygen-limited conditions (**Fig. 5e**). These findings could have implications for targeted therapies and the development of prognostic markers based on *FLAD1* expression levels and mitochondrial metabolic signatures in cancer.

### Design and screening of novel inhibitors of FLAD1

The essential role of FLAD1 in FAD biosynthesis positions it as a critical node in the metabolic adaptation of cancer cells to a low-oxygen environment. We have identified lumiflavin, a riboflavin analog, inhibits the cellular uptake of riboflavin and selectively inhibits the growth of hypoxic tumor cells (**Extended Data Fig. 7d**).

Given the absence of existing FLAD1 inhibitors, we utilized virtual screening to identify hit compounds targeting the FAD synthesis domain of FLAD1. We designed a workflow to computationally screen and optimize potential compounds with inhibitory effects against FLAD1 (**Fig. 6a**). As no protein structure of human FLAD1 has been resolved to date, we performed a homologous sequence search and modelled the protein using the structure of FAD synthase of *Saccharomyces cerevisiae* as a template (PDB code: 2WSI) and predicted the binding mode of FLAD1 and FAD (**Supplementary Table 3**, **Extended Data Fig. 11a**). A recombinant protein expression system was established to purify truncated FLAD1 protein *in vitro* (**Extended Data Fig. 11b**). Isothermal titration calorimetry (ITC) validated that this truncated protein retains a strong affinity for its substrate, FMN, indicating its potential utility in the development of a screening system for protein inhibitors (**Extended Data Fig. 11c**). Exploiting the differential fluorescent properties of FLAD1 substrate FMN and the product FAD, we developed an assay system for evaluating FLAD1-targeted inhibitory activity (**Extended Data Fig. 11d**). Virtual screening process generated six compounds for further experimental validation, among which, the compound C004 exhibited the strongest activity (**Supplementary Table 4**). C004 was predicted to form hydrogen bonds with N404, C409 and G487 while its benzofuran group formed Cation-Pi interaction with R513 and Pi-Pi interaction with W508, respectively. To further improve the bioactivity of C004, its derivatives were designed, synthesized and assessed for their inhibitory effect against FLAD1. Taking the carboxyl hydrazide group of C004 as the center, we modified the benzofuran group (R1), the methoxybenzene group, and the ethyl 2-hydroxyacetate group connected to methoxybenzene (R2), predicting the binding affinities of each modified molecules to FLAD1 (**Fig. 6a**, **Supplementary Table 5, 6, 7**). Based on the analysis of the binding modes, we synthesized several derivatives and tested them experimentally for their potential to inhibit FAD synthetic activity. Meanwhile, taking the carboxyl hydrazide group as the query, we conducted a substructure search for identifying compounds containing this important functional group and performed molecular docking on FLAD1 with these compounds. In total, fifteen compounds were screened for further validation (**Supplementary Table 8**). Consequently, we identified several compounds that inhibited FAD synthetic activity, among which compound C4-31 effectively inhibited the activity of recombinant FLAD1 protein. The main interactions of C4-31 are consistent with C004, including hydrogen bonds formed with N404, C409 and G487; however, stronger Pi-Pi interaction may be formed due to the substitution of the naphthalene group at R1 position.

**Fig. 6.**
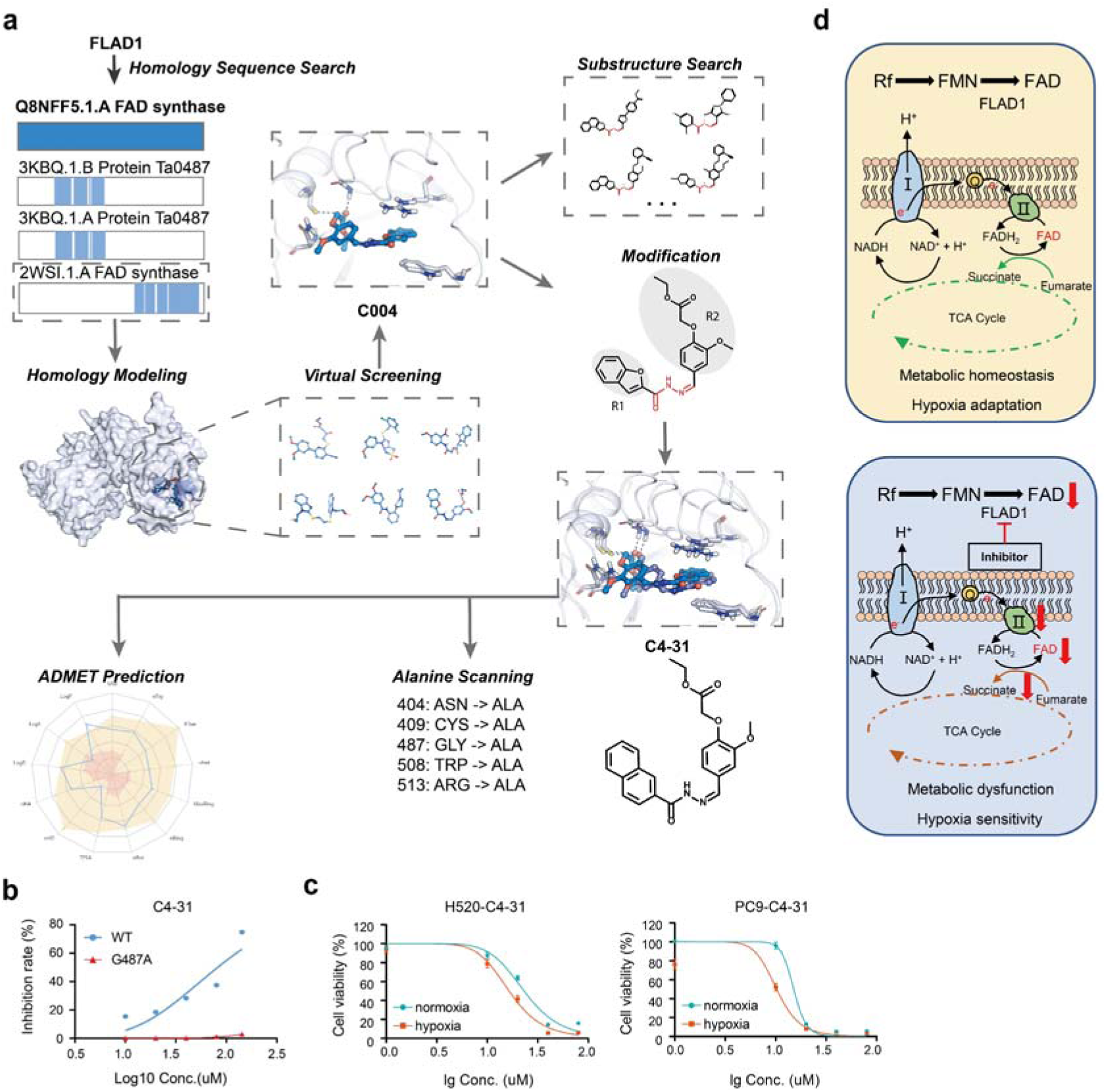
Discovery, optimization, and functional characterization of a novel FLAD1 inhibitor. **a,** Computational workflow of the discovery and optimization of compounds that inhibit FLAD1. **b**, Half maximal inhibitory concentration (IC50) of compound C4-31 against recombinant FLAD1 protein and its G487A mutant variant. **c**, IC50 of compound C4-31 on cells under normoxic and hypoxic conditions. **d**, The model illustrates the mechanism by which FLAD1 knockout or inhibition makes cells unable to tolerate hypoxic conditions. FLAD1 is involved in the intracellular synthesis of FAD. Under hypoxic conditions, oxygen cannot serve as the final electron acceptor in the electron transport chain (ETC), preventing the transfer of electrons. The activity of complex II of the ETC is reversed, primarily manifested as fumarate reductase activity. Electrons are smoothly introduced into fumarate, thus maintaining the flow of electrons within the ETC under low oxygen conditions. Inhibition or knockout of FLAD1 compromises the intracellular FAD pool, thereby impairing the activity of complex II, which relies on FAD as a cofactor, leading to dysfunction in the metabolic pathways coupled to the ETC and subsequent mitochondrial dysfunction.

To further confirm that the binding site of C4-31 is consistent with our prediction, we performed alanine scanning on the complex structure of FLAD1 and C4-31 to predict amino acid mutations that would strongly affect the stability of the protein and the binding affinity of the compound to the protein (**Supplementary Table 9**). G487, W508 and R513 were selected for subsequent experiments. Although G487 was predicted to have minimal effect on the binding affinity between the compound and FLAD1, it was also expected to have negligible impact on protein stability; thus, the protein activity would not be significantly affected (**Extended Data Fig. 11d**). Mutations of W508 and R513 were predicted to significantly reduce binding affinity and affect protein stability, which would negatively impact protein activity. Subsequent experimental results showed that the protein activity of these two mutant proteins was low, while the inhibitory effect of C4-31 against the FLAD1 G487A mutant decreased significantly (**Fig. 6b**), which agrees well with our computational simulation results and confirms that the binding of the compound is consistent with our predictions. We also performed an ADMET prediction for C4-31 to evaluate its pharmacological properties, and it was predicted to comply with Lipinski rule. Pan assay interference compounds (PAINS) are compounds that often result in false positive outcomes in high-throughput screening. PAINS tend to engage in non-specific interactions with many biological targets, rather than specifically affecting a desired target (Baell & Walters, 2014). Fortunately, C4-31 was predicted not to be a PAINS molecule. The predicted properties of C4-31 are listed in **Supplementary Table 10**. These predictions suggest that our compounds are drug-like molecules that deserve to be further developed.

### Targeted inhibition of FLAD1 selectively inhibits hypoxic tumors

Treatment of cells with the compound results in decreased intracellular FAD levels, demonstrating its ability to selectively target and inhibit intracellular FAD synthesis (**Extended Data Fig. 11e**). Testing the compound on FLAD1 wild-type and knockout cells, we observed that *FLAD1* knockout cells exhibited reduced sensitivity to the compound (**Extended Data Fig. 11f**), while the compound showed selective inhibition of hypoxic tumor cells (**Fig. 6c**). Here, we delineate gene *FLAD1*, which is integral to tumor hypoxia adaptation. Loss or inhibition of FLAD1 leads to decreased intracellular FAD levels, resulting in compromised mitochondrial complex II activity and an imbalance of succinate and fumarate. This culminates in mitochondrial metabolic dysregulation, rendering the cells incapable of tolerating hypoxic conditions (**Fig. 6d**).

## Discussion

Here, we successfully developed a deep learning model “DepFormer” to predict the dependent genes for each specific cell. These *in silico* predicted genes are significantly enriched in the hits identified by CRISPR screening in cells, suggesting that these genes are indeed required for the survival of cells. The constructed “DepFormer” model can be applied to identify dependent genes for cells of different status, including the dependent genes for cells under hypoxia.

Metabolic adaptation of tumor cells to hypoxic environment is still not completely understood, and therapeutic targets for hypoxic tumors are still lacking. Here, we performed systematic metabolic network comparison between hypoxic and non-hypoxic tumor cells, and identified oxidative phosphorylation (OXPHOS) as the most distinct metabolic pathway. Through the constructed “DepFormer” deep learning model, OXPHOS was identified as the primary dependent metabolic pathway for the survival of hypoxic tumor cells. Despite the established Warburg effect, which suggests that cancer cells prefer glycolysis regardless of oxygen availability, recent studies have begun to delineate a more nuanced relationship between cancer cell metabolism and oxygen levels, indicating that metabolic adaptation is context-dependent (Fendt *et al*, 2020; Jia *et al*, 2019; Palm, 2021). Our findings are aligned with this emerging paradigm. Our deep learning model-driven research has elucidated a fascinating direct link between hypoxia and the reliance on OXPHOS, specifically mitochondria ETC complex II, in the metabolic adaptation of hypoxic tumor cells. Contrary to the conventional belief, many cancers retain robust OXPHOS capability which can be up-regulated in certain malignancies like leukemia, lymphoma, and some solid tumors, including pancreatic ductal adenocarcinoma(Weinberg & Chandel, 2015; de Beauchamp *et al*, 2022; Zhang *et al*, 2019; Masoud *et al*, 2020). This revelation has propelled the exploration of OXPHOS inhibitors as therapeutic agents. Targeting OXPHOS may represent an effective strategy for cancers that heavily rely on mitochondrial respiration.

Here we identify *FLAD1*, a critical gene associated with the OXPHOS pathway, mediated mitochondria metabolic reprogramming that is required for tumor hypoxia adaptation. *FLAD1* emerged with a high-ranking perturbation score in our model, exhibiting significantly differential expression in tumors compared to normal tissues. This suggests that FLAD1 plays a pivotal role in the cellular response to hypoxic conditions often encountered within the tumor microenvironment. Subsequent experimental validations confirmed that *FLAD1* depletion specifically suppress the proliferation of cells under hypoxia. Conversely, up-regulation of FLAD1 was found to enhance tumor cell adaptation to hypoxia. FLAD1 is known to catalyze the synthesis of FAD in mammalian cells, and mutations in this gene have been identified as the etiological factor in approximately 2% of cases of mitochondrial FAD related metabolic disorder MADD, an autosomal recessive disease(Prasun, 1993; Lupica *et al*, 2022). The precursor of FAD, riboflavin, has been reported in some studies to be associated with hypoxia at high altitudes(Wickson & Morgan, 1946). Animals deficient in riboflavin are unable to effectively cope with high-altitude hypoxia; however, upon administration of riboflavin to riboflavin-deficient mice, they rapidly become better equipped to deal with high-altitude hypoxia(Riboflavin and the Stress Response, 1955; Liu *et al*, 2010). These previous observations are in line with the tumor hypoxia adaptation function of FLAD1 reported in this study. FAD, as a cofactor for various flavoproteins, is involved in many critical biological processes, including the citric acid cycle, β-oxidation, amino acid degradation, and the demethylation activity of LSD1(Lienhart *et al*, 2013; Becker *et al*, 2011; Yang *et al*, 2017; Barbara *et al*, 2010). Here, we uncover a novel function of FAD in tumor hypoxia adaptation.

FLAD1 functionally associates closely with mitochondrial ETC complex II. As part of OXPHOS, complex II, also known as succinate dehydrogenase (SDH), occupies a unique niche at the interface of the TCA cycle and the ETC. Three primary subunits within complex II, namely SDHB, SDHD, and SDHA, also exhibit elevated perturbation scores in our deep learning model. Mitochondria ETC complex II could exhibit reverse activity under hypoxic conditions, acting as a fumarate reductase that reduces fumarate to succinate, and this reverse activity has been observed in bacteria, yeast, animals, and specific parasites, reflecting a metabolic flexibility evolved by organisms to adapt to hypoxia, which may serve as a potential therapeutic target (Kita *et al*, 2002; Lemire & Oyedotun, 2002; Maklashina *et al*, 1998; Spinelli *et al*, 2021).

Our study, identifies *FLAD1* as a key gene in cellular adaptation to hypoxia, opening a new avenue for the development of cancer metabolic therapies and positioning this gene as a target for selective cancer treatments. Through molecular docking, we developed and validated the first FLAD1 specific inhibitors. Recently, a study reported that Chicago sky blue 6B can inhibit FLAD1 activity(Nisco *et al*, 2023). Chicago sky blue 6B was originally identified as a compound with antimicrobial activity through targeting the FADS of Corynebacterium ammoniagenes (CaFADS) (Sebastián *et al*, 2018). Prokaryotic FADS is functionally and structurally different from human FLAD1. CaFADS catalyzes the reaction from riboflavin to FAD, while human FLAD1 catalyzes the formation of FAD from FMN. After careful evaluations, we observed that Chicago sky blue 6B may not be a specific FLAD1 inhibitor for the following reasons: first, the compound has been predicted to belong to pan-assay interference compounds (PAINS) by ADMET2.0 (https://admetmesh.scbdd.com/) due to the presence of an azo group. We experimentally confirmed that Chicago sky blue 6B can inactivate proteins non-specifically. Second, under the ADMET prediction by ADEMT2.0 (Xiong *et al*, 2021), the compound was predicted to violate the Lipinski’s Rule of Five, indicating it is a non-drug-like molecule.

Furthermore, Chicago Sky Blue 6B contains a highly polar sulfate group, which may lead to the formation of non-specific ionic bonds between the compound and FLAD1 (**Supplementary Table 10**). The IC50 of our developed inhibitor exceed 1 uM, we are still in the process of optimizing this inhibitor for clinical application. It is also imperative to explore whether the inhibitory effects of FLAD1 can be synergistically combined with other therapeutic modalities. Moreover, further investigation into the role of FLAD1 within the tumor microenvironment and its interactions with other cellular stress responses is warranted.

In summary, the construction of deep learning model for predicting the hypoxia-dependent genes could inspire similar research aimed at identifying therapeutic targets in cancer cells. Furthermore, our study not only highlights the pivotal role of FLAD1 in the metabolism of cancer cells, particularly under hypoxia, but also proposes a feasible therapeutic target exploiting the metabolic vulnerability of cancer.

## Acknowledgments

We thank ShanghaiTech University High Performance Computing Public Service Platform for computing services. We thank multi-omics facility, molecular and cell biology core facility of ShanghaiTech University for technical help. This work is supported by National Natural Science Foundation of China (82373149), Shanghai Science and Technology Commission (24J22800700), open project fund of the National Health Commission’s key laboratory of individualized diagnosis and treatment of nasopharyngeal cancer (2023NPCCK02), cross disciplinary Research Fund of Shanghai Ninth People’s Hospital, Shanghai JiaoTong University School of Medicine (JYJC202227) and startup funding from ShanghaiTech University.

## Author Contributions

X.Z. performed all cell line and mouse model related biochemical and metabolomics experiments, and drafted the manuscript under the supervision of X.-S. L. T.W. performed metabolic network analysis and constructed the deep learning model for predicting metabolic dependent genes and pathways. S.W. performed molecular docking, designed FLAD1 inhibitor, and wrote the text related to FLAD1 inhibitor design under the supervision of F.B. Y.C. synthesized the FLAD1 inhibitors under the supervision of J.C. T.L., Y.Z., J.C., K.D., Z.H., J.Y., C.X., L.L. participated in cancer genome analysis and cell line related experiments. G.F., D.X., X.L. participated in critical project discussion. X.X. provided critical materials for this study and participated in critical project discussion and supervision. X.-S. L. conceived this study, identified FLAD1, designed the experiments, supervised this study and wrote the manuscript with input from X.Z. and T.W.

## Declaration of interests

The authors declare that they have no conflict of interest.

## Methods

### Dataset description for model construction

The RNA expression data from The Cancer Genome Atlas (TCGA) is sourced from the UCSC Xena platform, available at https://xenabrowser.net/. The abbreviations for the types of tumors analyzed and their corresponding full names are provided in **Supplementary Table 1**. The hypoxia scores for TCGA tumor patients are based on the study by Bhandari et al(Bhandari *et al*, 2019).Tumor specific genome-scale metabolic (GEMs) were obtained from Francesco Gatto et al(Gatto *et al*, 2020), which contains 4849 TCGA tumor samples. Lung cancer single cell data were downloaded from the studies of Prazanowska, K.H. et al. and Qian, J. et al(Prazanowska & Lim, 2023; Qian *et al*, 2020), with the former used as a training set and the latter as a validation set. Protein-protein interaction data was obtained from the STRING database using the R package rbioapi. The hypoxia-related gene set was obtained from Yanru Zhang et al(Zhang *et al*, 2023), and the cell death-related gene set was obtained from the study of Yutian Zou et al(Zou *et al*, 2022). The CRISPR-Cas9 screen data of 81 lung cancer cell lines was downloaded from Depmap project (23Q4 version). The dependency probabilities value is between 0 and 1. Genes with a dependency score > 0.8 are marked as cell dependent, genes that are dependent in more than 20 cell lines are marked as CRISPR hit, and other genes are marked as CRISPR not-hit.

### Enzyme Network Analysis

Sample specific GEMs were downloaded in mat format and converted to XML format by Matlab. Then we applied the R package Met2Graph to extract enzyme network from GEMs(Granata *et al*, 2022). In this network, enzymes are the nodes connected by edges represented by metabolites. Two enzymes are connected if they catalyze two reactions which produce or consume a specific metabolite.

We use Graph2Vec(Narayanan *et al*, 2017), an algorithm designed to learn vector representations for whole graphs, to compute the embedding vector for each sample’s enzyme network. Then we use cosine similarity to calculate the similarity between pairs of enzyme networks. Based on the Buffa hypoxia score(Buffa *et al*, 2010), samples are divided into hypoxic (score greater than the 75th percentile) and non-hypoxic groups (score less than the 25th quartile). We compared the differences in network similarity between samples with the same hypoxia status and those with different hypoxia statuses. We also calculated metrics to measure the importance of gene nodes in the enzyme network, including degree centrality, which indicates the number of connections a gene node has with every other gene; betweenness centrality, quantifying the number of times a gene node appears on the shortest path between two other nodes; eigenvector centrality which quantifies a node’s influence in the network based on its connections to other high-scoring gene nodes and closeness centrality, which calculates the length of the shortest path between a gene and all other genes in the network. These metrics were calculated by R package igraph. We retrieved the gene-wise vector of centrality measures of each sample specific enzyme network and computed the euclidean distance between these vectors of pairwise samples. Then the differences in this network distance between samples with the same hypoxia status and those with different hypoxia statuses were calculated.

For each gene, we calculated the difference in the previously mentioned centrality metrics between hypoxic and non-hypoxic tumors. We defined a gene as havin a significant up-regulated metric if it had an FDR value less than 0.05 (wilcoxon rank-sum test) and the median of the centrality metric in hypoxic tumor samples was greater than in non-hypoxic tumor samples. We filtered for genes with at least one significantly up-regulated centrality metric and used these genes to perform over representation enrichment analysis of metabolic pathways.

### *In Silico* Perturbation Analysis

The raw expression counts matrix for lung cancer single cell data was downloaded from the corresponding study and processed using the R package Seurat. To only retain high quality data, we removed all cells that have fewer than 250 genes with mapped reads and contain more than 15% of mitochondrial specific reads. Then, we used TCfinder (https://github.com/XSLiuLab/TCfinder) to predict cancer cells and retained only the cancer cells for downstream analysis. AUCell was used to calculate activity score of hypoxia gene set and cell death related gene set for each cell. Gaussian mixture model (GMM) was used to assign cells into high- and low-score group based on cells’ activity scores. The high or low-score groups were named as hypoxia/dying state and non-hypoxia/non-dying state respectively.

The data from K.H. et al was used as training data(Prazanowska & Lim, 2023), and 30% of the training data was applied to monitor training process and tune hyper-parameters. The hyperparameter tuning process is implemented by the Ray Tune framework and used AUC score as the evaluation metric. After hyper-parameter tuning, we set following hyper-parameters: 3.152009e-04 for learning rate, cosine for learning rate scheduler, 635.01 for warmup steps, 0.279 for weight decay and 24 for batch size. The fine-tuned model was independently validated on data sets from Qian, J. et al. In order to do *in silico* perturb, we merged the training set and test set to increase the sample size. *In silico* perturbation was achieved by removing the given gene from the rank value encoding of the given single-cell transcriptome and quantifying the cosine similarity between the original and perturbed cell embeddings to determine the predicted deleterious impact of deleting that gene in that cell. This impact was compared with the random distribution drawn from the other genes to calculate p value and corresponding FDR.

We first performed *in silico* perturbation on all single cells, identified genes whose deletion could significantly shift cell embedding from the non-dying state to the dying state, and compared these genes with the dependent genes obtained by CRIPSR. The *in silico* perturbation hit genes were defined as the effect value obtained by deleting the gene was > 0, and the FDR was < 0.05 (the effect value refers to the magnitude of cell embedding shift from the non-dying state to the dying state after deleting the specific gene). We extracted genes which interact with OXPHOS pathway genes from the STRING database to identify OXPHOS-associated genes and construct a protein-protein interaction network.

Hypoxia specific dependent genes were obtained by comparing *in silico* perturbation results in hypoxic cells and non-hypoxic cells. We filtered the results of *in silico* perturbation by the following criteria: 1. The effect value obtained by deleting the gene under hypoxia state was > 0, and the FDR was < 0.05; 2. Based on the TCGA dataset for LUAD and LUSC, we identified genes that are significantly upregulated in lung cancer compared to normal tissues (log2 fold change > 0 and adjusted p-value < 0.05); 3. We analyzed the TCGA dataset to identify genes that are significantly upregulated in hypoxic lung cancer tissues compared to non-hypoxic lung cancer tissues (log2 fold change > 0 and adjusted p-value < 0.05); 4. We excluded pan-essential genes from our analysis, utilizing data obtained from the DepMap Portal (24Q2). For each gene we defined perturbation scores: Perturbation Score = (-log10(Perturbation-FDR)) * Shift. The fold change indicates the selectivity of gene perturbations in hypoxic tumors, calculated as the perturbation score under hypoxic conditions divided by the perturbation score under normoxic conditions.

### Analysis of FLAD1 mRNA expression characteristics

We utilized the UCSCXenaTools R package(Wang & Liu, 2019) to obtain the FLAD1 mRNA levels (log2(FPKM+1)) data for 20 types of tumors as recorded in **Supplementary Table 1**. For comparison between tumor and non-tumor samples, we initially classified them into two groups based on their TCGA barcode. The differences between groups were analyzed using the Wilcoxon rank-sum test. The first row above the x-axis indicates the number of non-tumor samples, while the second row indicates the number of tumor samples.

To analyze the correlation between FLAD1 and hypoxia, we calculated the Pearson correlation coefficient between each gene in tumor samples and the Buffa hypoxia score, after segregating by cancer type. Subsequently, the correlation of each gene in tumors with the Buffa hypoxia score was visualized using box plots, with red diamond markers highlighting the correlation of FLAD1 with the hypoxia score. The sample counts are annotated above the x-axis.

### Analysis of FLAD1 copy number features

We utilized the UCSCXenaTools R package to retrieve the FLAD1 GISTIC copy number data for the 20 types of tumors as documented in **Supplementary Table 1**. Based on the hypoxia levels indicated by the Buffa hypoxia score, we selected the top 30% as the hypoxic group and the bottom 30% as the non-hypoxic group among the tumor samples. We then separately visualized the FLAD1 copy number status for LUAD and Pancancer. Subsequently, we employed Fisher’s exact tests to compare the significance between the hypoxic and non-hypoxic groups

### Cell culture

In our investigation, all cell lines were utilized within 20 passages or less than four months post-resuscitation to ensure their optimal state, coupled with regular assessments for mycoplasma contamination. Four distinct cell lines—H520, H1299, A549, and PC9—were cultured in RPMI 1640 medium (11875093, Gibco), supplemented with 10% fetal bovine serum (FBS) (10091148, Gibco) and 1% penicillin-streptomycin (15140122, Gibco). HEK-293t cells were maintained in DMEM (11995065, Gibco), similarly enriched with 10% FBS and 1% penicillin-streptomycin (15140122, Gibco). All cell lines were incubated in a controlled environment at 37°C with a 5% CO2 within a cell culture incubator.

### Establishment of FLAD1 knockout cancer cell lines

For the purpose of targeting the FLAD1 gene, two sgRNAs were designed using the Benchling platform (www.Benchling.com): sgRNA1: AAACAGCTCATCTCTCTCTC and sgRNA2: CAGAAGTAGGGGTCGGGCGT. The plasmid pUC57kan-T7-gRNA-U6 V2 (115520, Addgene) was employed as the PCR template to amplify the sgRNA1-U6-sgRNA2 PCR product. The pGL3-U6-sgRNA-ccdB-EF1a-puro plasmid was digested with BsmBI enzyme (R0739S, NEB), followed by ligation using T4 Ligase (M0202, NEB) to generate a plasmid harboring both sgRNAs. The ligation product was transformed into DH5α competent cells, which were incubated on ice for 30 minutes, subjected to a heat shock at 42°C for 90 seconds, and then rapidly cooled on ice for 2 minutes. The cell-plasmid mixture was then combined with LB medium and incubated at 37°C with gentle shaking (150 rpm) for 1 hour. Transformed cells were collected by centrifugation and plated on appropriate agar plates. Colonies were selected and sequenced to confirm the accurate insertion of sgRNAs into the plasmid, yielding the pGL3-2U6-2sgRNA-EF1a-Puromycin (pGL3-sgRNA) construct.

H1299, H520, and PC9 cells were cultured in 6-well plates to achieve 70-90% confluency. For transfection, 2µg of pGL3-sgRNA and 2µg of pST1374-NLS-flag-linker-Cas9-blasticidin plasmids were each diluted in 250µL of serum-free Opti-MEM medium following the manufacturer’s protocol for Lipofectamine 2000. The mixture was then added to the cells for incubation. After 48 hours, the cells were treated with puromycin (10mg/mL, ST551-10mg, Beyotime) and blasticidin (60218ES10, YEASEN). Control groups were simultaneously treated with corresponding concentrations of puromycin and blasticidin S for monitoring; when these control cells were completely killed, the next step was carried out.

Remaining transfected cells were digested and subjected to single-cell sorting into 96-well plates using the BD FACSARia™ III. Individual cells were allowed to proliferate in the wells until they reached sufficient numbers. DNA was extracted from the expanded single cells using a Quick-DNA extraction kit (QE0905T, Lucigen). PCR screening for FLAD1 knockout was performed using primers designed for the purpose (Primer sequences: AAGGGGTGAGGTCTCCT and GGTGATAAACCTCTCCTCCA). Successful knockouts should display a band of approximately 300bp, while wild-type cells will have a band of around 500bp. Proteins were extracted from amplified, validated knockout cells using proteinase inhibitors and RIPA lysis buffer. Protein concentrations were quantified using the Bradford Protein Assay Kit. Protein expression was analyzed by western blot using specific antibodies.

### Establishment of FLAD1 overexpression cancer cell lines

The full-length sequence of FLAD1 was cloned from cDNA of the 293T cell line using PCR (Primer sequences: CGGAATTCGCCACCATGGGTTGGGATTTGGGAACAC and CGCGGATCCTTTCATGTGCGGGAGTTCCGCTCCTCTTCT). The pCDH-puro plasmid was digested with restriction endonucleases EcoRI (R0101S, NEB) and BamH1 (R0136S, NEB). The ligation reaction was performed using T4 DNA ligase. The construct was transformed into DH5α competent cells. Colonies were screened and the plasmid with the correct sequencing results was identified as pCDH-CMV-FLAD1-puro.

For virus production, 6µg of the pCDH-CMV-FLAD1-puro plasmid was mixed with the packaging plasmids pMD2.g (1.5µg) and psPAX2 (4.5µg) in OPTI-MEM (Gibco) to form the DNA mixture. The control group employed the pCDH-CMV-puro plasmid. The EZ-Trans transfection reagent (LIFE iLAB BIO, Shanghai, China) was mixed with the DNA mixture according to the transfection reagent’s protocol and incubated at room temperature for 20 minutes. The DNA-EZ mixture was then added to 293T cells at approximately 70% confluency in a 10 cm cell culture dish. After 7 hours, the cells were refreshed with new culture medium. The viral supernatant was collected 48 hours later, filtered through a 0.45µm filter, and temporarily stored at 4°C for future use.

A549 and PC9 cells were seeded in 6-well plates the night before at approximately 20% confluency. The culture medium containing viral particles was added, and the cells were co-cultured for 48 hours. Afterward, the cells were refreshed with new culture medium and cultured overnight. Puromycin (10mg/mL, ST551-10mg, Beyotime) was diluted 10,000-fold to select cells. Subsequently, the expression levels of FLAD1 in the cells were validated using qPCR and Western blot (WB).

### RNA extraction and quantitative real-time PCR

Cells were cultured in 6 cm dishes and total RNA was extracted using the HiPure Total RNA Mini Kit (Magen, P1112-03). Subsequently, cDNA was synthesized from 1μg of RNA via reverse transcription using the HiScript III 1st Strand cDNA Synthesis Kit (Vazyme, R312-01). Real-time quantitative PCR (qPCR) was performed using 20ng of cDNA with the SYBR Green qPCR Master Mix (Vazyme, Q331-02). Primers used are listed in the **Supplementary Table 11**. Accumulation of fluorescent PCR products during the PCR program (initial denaturation at 95°C for 5 minutes, followed by 40 cycles of 95°C for 10 seconds, and 60°C for 30 seconds) was monitored using the QuantStudio 7 system (Thermo Scientific). Gene expression was quantified relative to the housekeeping gene beta-actin using the ΔΔCt method.

### Western blotting

Cells in 6-well plates were digested with trypsin and subsequently washed three times with 1X PBS. Lysis was performed with RIPA lysis buffer, which was combined with a protease inhibitor cocktail in a 100:1 ratio. Proteins were resolved by electrophoresis on a 12% SDS-PAGE gel. The separated proteins were then transferred to a polyvinylidene fluoride (PVDF) membrane. Western blot analysis was conducted using anti-FLAD1 (ab95312, abcam) and anti-tubulin (ab59680, abcam). Chemiluminescent detection was achieved using horseradish peroxidase-conjugated secondary antibodies coupled with Western ECL reagent (ED0016-B, Sparkjade).

### Cell proliferation assay

Cells in the logarithmic growth phase from each experimental group were prepared as a single-cell suspension and seeded into a 96-well plate at a density of 4,000 cells per well. Each group was represented by triplicate to sextuplicate wells. Two days post-seeding, 10 μL of CCK-8 solution (C0005, Targetmol) was added to each well, including blank control wells containing only culture medium and CCK-8 solution. After incubating for one hour, the absorbance of each well was measured at 450 nm using a microplate reader and the optical density (OD) values were duly recorded.

### Mouse models

All animal experiments were approved by the Institutional Animal Care and Use Committee (IACUC) of ShanghaiTech University (approval number: 20200918002). Female BALB/c nude mice, 4 weeks of age, were obtained from Shanghai lingchang Animal Co., Ltd. (Shanghai, China), and housed in microisolator cages. Tumor cells (4 × 10^6 cells in 100 µL PBS) were subcutaneously injected into the flanks of the mice (n=7 per group). Tumor volumes were measured every three days and calculated using the formula: V = (L × W^2)/2, where V is the tumor volume, W is the tumor width, and L is the tumor length. Once the tumors reached a predetermined volume, the mice were euthanized, and the tumors were excised and weighed.

### Immunofluorescence imaging

Cells were cultured overnight on 20 mm glass-bottom dishes. They were fixed with 4% paraformaldehyde for 20 minutes and permeabilized with 0.5% Triton X-100 in PBS, followed by blocking with 1% bovine serum albumin (BSA) in PBS. After washing with PBS, cells were incubated with anti-FLAD1 (ab95312, Abcam) in 1% BSA in PBS at 4°C overnight. As a negative control, cells were incubated under the same conditions with 1% BSA in PBS. Following washes, cells were relabeled with an appropriately conjugated Alexa Fluor 488 secondary antibody at 37°C for 1 hour. The nuclei and mitochondria were stained using 4’,6-diamidino-2-phenylindole (DAPI, D9542, Sigma) and MitoTracker® Red CMXRos (40741ES50, YEASEN), respectively, following the protocols provided in the respective manuals. Cellular analysis was conducted using a Leica TCS SP8 confocal laser-scanning microscope (Leica Microsystems, Wetzlar, Germany) equipped with a 63× objective lens.

### Oxygen consumption rate measurement

Oxygen consumption rate (OCR) was measured using the Seahorse XFp Extracellular Flux Analyzer (Seahorse Bioscience) in accordance with protocols provided by the manufacturer. The Seahorse XFp Cell Mito Stress Test Kit (103010-100, Agilent Technologies) was utilized to assess OCR. Briefly, cells were harvested and seeded at 10,000 cells per well in a Seahorse XFp cell culture microplate with complete growth medium and cultured overnight. The following day, the medium was replaced with XF Base Medium (Seahorse Bioscience) supplemented with 10 mM glucose, 1 mM pyruvate, and 2 mM glutamine. The plate with cultured cells was then loaded into the Seahorse XFp Analyzer to record baseline measurements. For the OCR measurements, oligomycin, FCCP, and a combination of rotenone/antimycin A (Rot/AA) were sequentially injected. The OCR results were normalized to the protein content of the cells. Data analysis was performed using the Seahorse XFp Wave software.

### Human Protein Atlas Analysis

Immunohistochemical (IHC) staining images for FLAD1 were obtained from the Human Protein Atlas (HPA) database (accessed June 2022; https://www.proteinatlas.org/), and quantification of positive cells was performed using ImageJ software (accessed June 2022; https://imagej.nih.gov/ij/) and an open-source plugin “IHC Profiler”(Varghese *et al*, 2014).

### Kaplan-Meier survival analysis

To investigate the association between FLAD1 expression and patient survival, we downloaded gene expression data and associated survival information for cancer patients from the UCSC Xena platform. TCGA tumor samples were stratified into high (top 25%) and low (bottom 25%) FLAD1 expression groups. Survival outcomes for patients in these groups were evaluated and visualized using the survival package (version 3.5-7) and survminer package (version 0.4.9) in R. And the Cox proportional hazards regression model was used for hazard ratio estimations.

### Cell apoptosis assay

Cells were seeded at a density of 1×10^5 cells per well in a 6-well plate and incubated overnight. The following day, the cells were transferred to a hypoxia chamber to grow for an additional 72 hours before analysis. Cell staining was performed using propidium iodide (PI, 14289, Cayman) according to the manufacturer’s instructions, and the percentage of PI-positive cells was determined. Flow cytometric data was acquired on a CytoFLEX (Beckman Coulter) and analyzed using FlowJo software (version 10.01).

### Gene-set enrichment analysis

Gene expression data for multiple cancer types were retrieved from the UCSC Xena database. Patients within each cancer type were divided into two groups according to the median expression level of FLAD1: a high FLAD1 expression group and a low FLAD1 expression group. Differential gene expression analysis between the two groups was performed using the DESeq2 R package. Significantly upregulated genes in the high FLAD1 expression group for lung cancer were selected and subjected to Gene Ontology (GO) analysis using the clusterProfiler package (version 4.10) in R. Gene sets for KEGG pathways were downloaded from The Molecular Signatures Database (MSigDB; c2.cp.kegg.v7.4). Gene set enrichment analysis (GSEA) of differentially expressed genes across various cancers was conducted using the fgsea package (version 1.28.0) in R. Based on the average normalized enrichment scores of multiple tumor types, we sorted them in descending order and selected the top 20 for display. Asterisks denote adjusted p-values below 0.05. We chose the KEGG OXPHOS gene set and then utilized the UCSCXenaTools R package to acquire the expression levels of these genes in different tumor samples from TCGA. To represent the overall activity level of this pathway in individual tumor samples, we normalized the OXPHOS genes within each tumor type separately and then calculated the average value of OXPHOS genes for each tumor sample. This average value measures the overall level of the OXPHOS pathway in the tumor sample. Subsequently, based on the median expression level of FLAD1 in different samples within the tumor type, we divided the tumor samples into two groups: FLAD1 high and FLAD1 low. The differences between groups were analyzed using the Wilcoxon rank-sum test.

### ROS and mitochondrial membrane potential measurement

Intracellular reactive oxygen species (ROS) levels were measured using 2’,7’-Dichlorodihydrofluorescein diacetate (H2DCFDA, H131224, Aladdin), according to the manufacturer’s protocol. Cells were seeded at a density of 2.0×10^5 cells per well in a 6-well plate. Following the treatment of all groups as mentioned, cells were washed thrice with PBS and then incubated with H2DCFDA for 30 minutes. Cells were subsequently washed with 1X PBS, collected in PBS, and the fluorescence was measured using a FACSCalibur flow cytometer (BD Biosciences). For the assessment of mitochondrial membrane potential, cells were incubated with JC-10 dye (40752ES60, YEASEN) following the manufacturer’s instructions. Briefly, cells were incubated with JC-10 staining solution at 37°C for 20 minutes. After staining, cells were gently washed twice with the JC-10 staining buffer provided in the kit to remove excess dye and minimize background fluorescence. The flow cytometer was equipped with the appropriate laser and filters to detect the JC-10 monomeric form (green fluorescence) and the aggregated form (red fluorescence). The ratio of red to green fluorescence was calculated for each sample, serving as an indicator of the mitochondrial membrane potential.

### Quantitative analysis of riboflavin metabolism in tumor samples

Gene expression data (counts) spanning multiple cancer types were retrieved from the UCSC Xena database, representing a comprehensive subset of TCGA. Hypoxic status was determined based on the Buffa hypoxia score. For each cancer type, the top 30% of samples with the highest Buffa scores were classified as hypoxic tumors, and the bottom 30% as non-hypoxic tumors. The differential expression analysis of the genes was performed using the DESeq2 R package. This analysis compared the expression of the riboflavin metabolism-related genes (*SLC52A1, SLC52A1, SLC52A1, RFK, FLAD1*) expression level between tumor and normal tissues, as well as between hypoxic and non-hypoxic tumor samples. Further, to compare riboflavin metabolism levels in tumors with varying degrees of hypoxia, each cancer type’s tumor samples were categorized into five groups according to their Buffa hypoxia scores. These groups were labeled as “Very Low”, “Low”, “Medium”, “High”, and “Very High” hypoxia levels. For each cancer type, expression levels of riboflavin-related genes were normalized to enable cross-sample comparison. Following normalization, the mean expression level of the riboflavin metabolism genes in each sample was calculated to serve as an indicator of the overall riboflavin metabolic activity in that sample. This approach allowed for a standardized assessment of the metabolic state across the spectrum of hypoxia within the cancer datasets.

### Lumiflavin treatment

PC9 and H520 cells were seeded in 96-well plates at a density of 5,000 cells per well and allowed to adhere overnight. Lumiflavin (BD136849, Bidepharm) was prepared as a stock solution in DMSO and filtered for sterilization. The stock solution was further diluted in culture medium to achieve final concentrations of 10 µM (PC9 cells) and 50 µM (H520 cells). Cells were treated with the respective concentrations of lumiflavin for 48 hours under hypoxic (1% O2) or normoxic (21% O2) conditions. Cell viability was assessed using CCK-8 following the manufacturer’s guidelines. Each experimental condition was performed in triplicate, and results were expressed as mean ± standard deviation.

### Complex II inhibitor

To investigate the survival differences under hypoxic conditions between FLAD1 wild-type and knockout cells upon treatment with complex II inhibitors, 3-nitropropionic acid (3-NPA, N5636, Sigma) and 2-thenoyltrifluoroacetone (TTFA, ab223880, Abcam), we cultured PC9 and H520 cells in both wild-type and knockout variants. Cells were placed in a controlled environment at normoxic (21% O2) and hypoxic (1% O2) conditions to mimic physiological and hypoxic states, respectively. For PC9 cells, we administered a treatment of 15 µM 3-NPA and 200 µM TTFA, while H520 cells received 30 µM 3-NPA and 150 µM TTFA. The cells were treated continuously for three days under both oxygen conditions. Following the treatment period, cell viability was quantified using the CCK-8 assay according to the manufacturer’s protocol. Cell viability percentage under hypoxic conditions was calculated relative to normoxic conditions.

### Mitochondrial complex II enzyme activity assay

Approximately 10^6 cells were harvested from a 6 cm cell culture dish and subsequently resuspended in 1X PBS, adjusting the protein concentration to 5.5 mg/mL. The activity of mitochondrial complex II was measured using the Complex II Enzyme Activity Microplate Assay Kit (ab109908, Abcam). Briefly, cells were lysed using a 1/10 volume of detergent solution followed by incubation on ice for 30 minutes. Centrifugation was performed at 12,000 x g for 20 minutes at 4°C, and the supernatant was collected. Fifty microliters of the sample were added to the assay plate, along with 50 µL of incubation solution to serve as a blank reference. After incubation at room temperature for 2 hours, each well was washed with 1X buffer and the liquid was discarded. Subsequently, 40 µL of phospholipids was added and incubated for an additional 30 minutes. Two hundred microliters of the activity solution were then added to each well. The optical density at 600 nm was measured using a spectrophotometer to quantify enzyme activity.

### Flavin adenine dinucleotide (FAD) assay

Intracellular FAD levels were quantified utilizing the Flavin Adenine Dinucleotide Assay Kit (ab204710, Abcam). The procedure began with the collection of approximately 10^6 cells. These cells were washed with cold PBS and resuspended in 400 µL of ice-cold FAD assay buffer. Then samples were centrifuged at 10,000 x g for 3 minutes at 4°C to remove insoluble material. The supernatant was carefully transferred to a fresh tube and kept on ice. Protein precipitation was accomplished using trichloroacetic acid (TCA, 80132618, SCR), as delineated here: a 100% (w/v) TCA solution was prepared alongside a 2M KOH neutralization solution. For protein quantification, 10 µL of the sample was used. Subsequently, 30 µL of 100% (w/v) TCA solution was added to 300 µL of FAD assay buffer, followed by vortexing. After 15 minutes on ice, samples were centrifuged at 4°C, 12,000 g for 5 minutes. The clear supernatant was then transferred to a new tube. Approximately 60 µL of KOH neutralization solution was added to adjust the pH to 6.5-8.0, followed by an additional centrifugation at 4°C, 12,000 g for 5 minutes, and the supernatant was again transferred. A 1/10 volume of the supernatant was used for the assay, with the volume being adjusted to 50 µL using assay buffer, before being added to the wells. The reaction mixture, comprising 46 µL of FAD assay buffer, 2 µL of oxired probe, and 2 µL of enzyme mix, was then added. Following thorough mixing, the optical density at 570 nm (OD570) was measured at 5-minute intervals to monitor the reaction progress.

### Stable-isotope tracing experiments

PC9 cells with wild-type FLAD1 and FLAD1 knockout were cultured in 6 cm cell culture dishes using RPMI 1640 medium supplemented with 10% dialyzed fetal bovine serum (dFBS) and 1% penicillin/streptomycin. When cells reached 80% confluence, the unlabelled 1640 medium was removed. The cells were washed twice with 4 mL 1x PBS. One set of cells (n=6) was switched to 1640 medium (no glucose, 11879020, Gibco) containing 2 g/L U-13C-glucose (D346444, Aladdin) and 10% dFBS, while the other set (n=6) was maintained in 1640 medium with unlabelled 2 g/L glucose. Cells were then cultured in a hypoxia chamber with 1% oxygen for 15 hours. Subsequently, the medium was discarded, and cells were washed twice with 1x PBS. The cell plates were placed on dry ice and 1 mL pre-cooled MeOH:H2O (4:1, v/v, at −80°C, Methanol, 1.06007.4008, Merck) mixture was added. The cells were incubated at −80°C for 40 minutes. The cells were scraped off the plate and transferred into 1.5 mL centrifuge tubes. The samples were vortexed for 1 minute and then centrifuged at 13,000 g for 10 minutes at 4°C and the supernatant was transferred to a vacuum concentrator for evaporation. Subsequently, 100 μL ACN:H2O (1:1, v/v) was added to the extract, followed by vortexing for 1 minute and centrifuging at 13,000 g for 10 minutes at 4°C to pellet insoluble materials. The clear supernatant was finally transferred to LC/MS vials for analysis.

### LC–MS analysis and data processing

For untargeted metabolomics analysis, LC–MS was performed using a Waters ACQUITY UPLC I-Class system (Waters) coupled with a Quadrupole Time-of-Flight mass spectrometer (TripleTOF 6600, SCIEX). The mobile phase consisted of 25 mM ammonium acetate and 25 mM ammonium hydroxide in 100% water (mobile phase A) and 100% acetonitrile (mobile phase B). Instrument parameters were optimized based on previously reported methods. The raw data acquired by the mass spectrometer in .wiff format were converted to .mzXML and .mgf files using MSConvert (version 3.0.23109). The generated MS1 peak lists and MS2 files were then submitted to MetDNA (http://metdna.zhulab.cn/) for metabolite annotation. Metabolites significantly downregulated in the FLAD1 knockout group were selected using the Wilcoxon non-parametric test and submitted to MetaboAnalyst (https://www.metaboanalyst.ca/) for pathway enrichment analysis. For targeted metabolomics, LC-MS analysis was conducted using an AB Sciex QTRAP 6500+ system coupled with a Nexera UHPLC LC-30A 30A system (Shimadzu). Quantification of metabolites in the results was performed using MultiQuant 3.0.1 (Sciex).

### Computational discovery of inhibitors against FLAD1

For the absence of 3D structure of human FLAD1, we first conducted a homology sequence search on the sequence of FLAD1 to search the template for structure modelling using SWISS-MODEL(Waterhouse *et al*, 2018). The structure of FAD synthase of Saccharomyces cerevisiae with the highest identity was chosen as a template for homology modelling (PDB code: 2WSI). The model was constructed by Robetta with all parameters set to default except for the template specified as 2WSI(Kim *et al*, 2004). The structure was prepared by Protein Preparation Workflow of Schrödinger2021-4 in which we filled in missing side chains and hydrogens and performed a restrained minimization(Sastry *et al*, 2013). The modelled structure was aligned with structure in 2WSI to confirm the binding site of FAD. Glide of Schrödinger was applied to dock FAD to FLAD1(Friesner *et al*, 2004). Grid file of FLAD1 was generated using the Receptor Grid Generation tool of Schrödinger, taking the location of FAD as the binding site(Repasky *et al*, 2007). Glide was then utilized to dock FLAD1 to compounds in Specs compound library. After visual inspection of the docking modes between compounds and FLAD1, the selected complexes of compounds with FLAD1 were optimized by Molecular Mechanics/Generalized Born Surface Area (MM/GBSA) of Schrödinger(Genheden & Ryde, 2015). The modified compounds were prepared by LigPrep tool of Schrödinger. They were also docked to FLAD1 by Glide and their complexes with FLAD1 were optimized by MM/GBSA as well. Substructure search of the carboxyl hydrazide group was performed on Specs website (https://www.specs.net/). The searched compounds were prepared and docked to FLAD1. Alanine scanning was conducted on the complex of FLAD1 and C4-31 utilizing residue scanning tool of Schrödinger with residues involved in the main interactions, including N404, C409, G487, W508 and R513, mutated to Alanine(Beard *et al*, 2013). Delta affinity and delta stability of the complex were observed. ADMET prediction was performed by ADMETlab2.0 to evaluate the pharmacological properties of compound C4-31(Xiong *et al*, 2021).

### Optimization of inhibitors

Our compound modification strategy is to fix the carboxyl hydrazide group while modifying the groups on both sides of it. At R1 position, we modified the benzofuran to groups like indonaphthene, quinoline, dihydroxyoteridine (**Supplementary Table 5**). Since that the aromatic ring at R1 position formed hydrophobic interactions with W508 and R513, the introduction of more aromatic rings may have a positive effect on the enhancement of interactions. For R2 position, we first modified the benzene ring to pyrrolidine, pyrrole, imidazole, pyrazole and furan (**Supplementary Table 6**). Furthermore, referring to the structure of FAD, we made an overall modification at R2 position. In addition, after computationally evaluating the bindings of FLAD and the molecules modified at R1 and R2 position, we tried to modify both positions to obtain better molecules (**Supplementary Table 7**).

### Expression, purification and identification of recombinant His-FLAD1

To amplify the cDNA of FLAD1 from 293T cell cDNA, forward primer 5′-GTGCCGCGCGGCAGCCATATGCCCAACGCTGTGGAGCA-3′ and reverse primer 5′-CCGCAAGCTTGTCGACGGTCATGTGCGGGAGTTCCGCTCCT-3′, both containing homology arms, were utilized. The pET28_T7_6xHis_lac plasmid was digested using NdeI and SacI restriction enzymes. Subsequently, the FLAD1 sequence was seamlessly cloned into the pET28 plasmid to generate the pET28-6xHis-FLAD1 construct. The pET28-6xHis-FLAD1 construct was transformed into Transetta (DE3) chemically competent cells (CD801-02, Transgen). We utilized overlap PCR to construct different FLAD1 mutant variants. Positive colonies were screened on LB agar plates containing 34 µg/mL chloramphenicol and 100 µg/mL kanamycin, followed by sequence verification. E. coli cells harboring the recombinant plasmid were inoculated into 10 mL LB medium supplemented with 34 µg/mL chloramphenicol and 100 µg/mL kanamycin and cultured overnight at 37°C with shaking at 160 rpm. The culture was then transferred at a 1:100 ratio to 0.5 L fresh LB medium supplemented with kanamycin and chloramphenicol and grown at 37°C until A600 reached 0.6–0.8. At that point, 0.4 mM IPTG was added to induce the expression of the 6-His-FLAD1 recombinant protein. Growth was continued overnight at 20°C. The next day, bacteria were harvested by centrifugation at 3000×g for 20 minutes at 4°C and the pellet was stored at −80°C. A lysis buffer containing 20mM Tris, 300mM NaCl, 5mM imidazole, and 1mM PMSF, with a pH adjusted to 7, was prepared. The E. coli pellet was resuspended in this buffer and lysed three times through an EmulsiFlex-C3 high-pressure homogenizer (Avestin) at 700 bar. Protein electrophoresis was performed on bacterial lysates with and without IPTG induction, and proteins were visualized using Coomassie Brilliant Blue R250 staining to preliminarily confirm protein expression. The homogenized solution was cleared of insoluble material by centrifugation at 10,000 rpm for 20 minutes at 4°C, and the supernatant was collected to obtain the recombinant protein solution. The recombinant protein was purified using Ni-NTA affinity chromatography (C600793, Sangon) following the manufacturer’s instructions. The wash buffer used in this process contained 20mM Tris, 300mM NaCl, and 5mM imidazole at pH 7. The elution buffer contained 20mM Tris, 300mM NaCl, and 500mM imidazole at pH 7.

After elution, further purification was performed using column chromatography (HiLoad 16\600 Superdex 200 pg, Cytiva) under buffer conditions of 20 mM HEPES, 200 mM NaCl, 1 mM TCEP, pH 7.5, to obtain FLAD1 protein with a purity >95%. Subsequently, 10% glycerol was added to the protein solution, aliquoted, and flash-frozen in liquid nitrogen.

### Isothermal calorimetry

To evaluate the binding parameters of FLAD1 protein with its substrate FMN, isothermal titration calorimetry (ITC) measurements were carried out using a MicroCal PEAQ-ITC system (Malvern). An ITC-compatible buffer (20mM Tris, 150mM NaCl, 5mM MgCl2, pH7.5) was prepared. The buffer of FLAD1 recombinant protein was exchanged using an Amicon Ultra-0.5 Centrifugal Filter Unit (UFC5003BK, Merck Millipore), and an FMN solution at a final concentration of 200 µM was prepared using the ITC buffer. A solution containing 10 µM of FLAD1 protein was placed in the sample cell, while a solution containing 200 µM FMN was placed in the reference cell. A typical experiment consisted of 13 injections, each of 2 µL, with an equilibration time of 150 seconds between injections. Data processing was performed using the analytical software provided by Malvern.

### Measurements of FAD synthesis rate

To measure the rates of FAD synthesis, exploiting the differential fluorescence characteristics of FAD relative to FMN, the methodology was optimized based on protocols reported in previous studies(Torchetti *et al*, 2011; Leone *et al*, 2018). Fluorescence kinetics were tracked at 37°C using a SpectraMax i3 Microplate Reader (Molecular Devices), with excitation at 450 nm and emission detection at 520 nm. FAD synthesis rate was expressed as nmol FAD/min/mg protein. For the enzyme activity assay, 1.35 µg of purified protein was incubated at 37°C in 50 mM Tris/HCl buffer at pH 7.5 containing 5 mM MgCl2, 5µM FMN, 100µM ATP, and the compound under investigation.

### IC50 measurement of FLAD1 inhibitors

For inhibitor treatments, cells were seeded into 96-well plates at a density of 4000 cells per well. After 24 hours, cells were treated with varying concentrations of FLAD1 inhibitors (1, 10, 20, 40, and 80 μM). Cells treated with 0.1% (v/v) DMSO served as the vehicle control. Post-treatment with the inhibitors for 48 hours, the efficacy was evaluated using the CCK-8 assay. Briefly, 10 μL of the CCK-8 solution was added to each well, and the plates were incubated for 1 hour at 37°C. The absorbance of each well was measured at 450 nm. Data from the CCK-8 assay were inputted into GraphPad Prism for IC50 calculations. The normalized response (viability %) was plotted against the log-transformed inhibitor concentrations. The IC50 was determined using a non-linear regression model (log(inhibitor) vs. normalized response - Variable slope). Each IC50 value represented the concentration of inhibitor required to decrease cell viability by 50% relative to the DMSO vehicle control. Experiments were performed in triplicate and repeated at least three times independently.

### Quantification and statistical analysis

For data analysis, R program (4.2.3), the GraphPad Prism Software (version 9.5, Vienna, Austria) and Microsoft Excel (Microsoft Office 2013 for Windows, Albuquerque, NM, USA) were used. Unless otherwise specified, results are presented as the mean ± standard error. Other statistical analysis was performed using the unpaired, two-tailed Student’s t-test and p < 0.05 was considered to indicate a statistically significant difference.

## Data and Code availability

The fine-tuned model, corresponding processed training and test data was archived in Zenodo (https://doi.org/10.5281/zenodo.10957878), codes required to reproduce the analysis in this manuscript are available at https://github.com/XSLiuLab/hypoxia_target, and analysis report are available online at https://xsliulab.github.io/hypoxia_target/.

## Supplementary information

**Extended Data Fig. 1.**
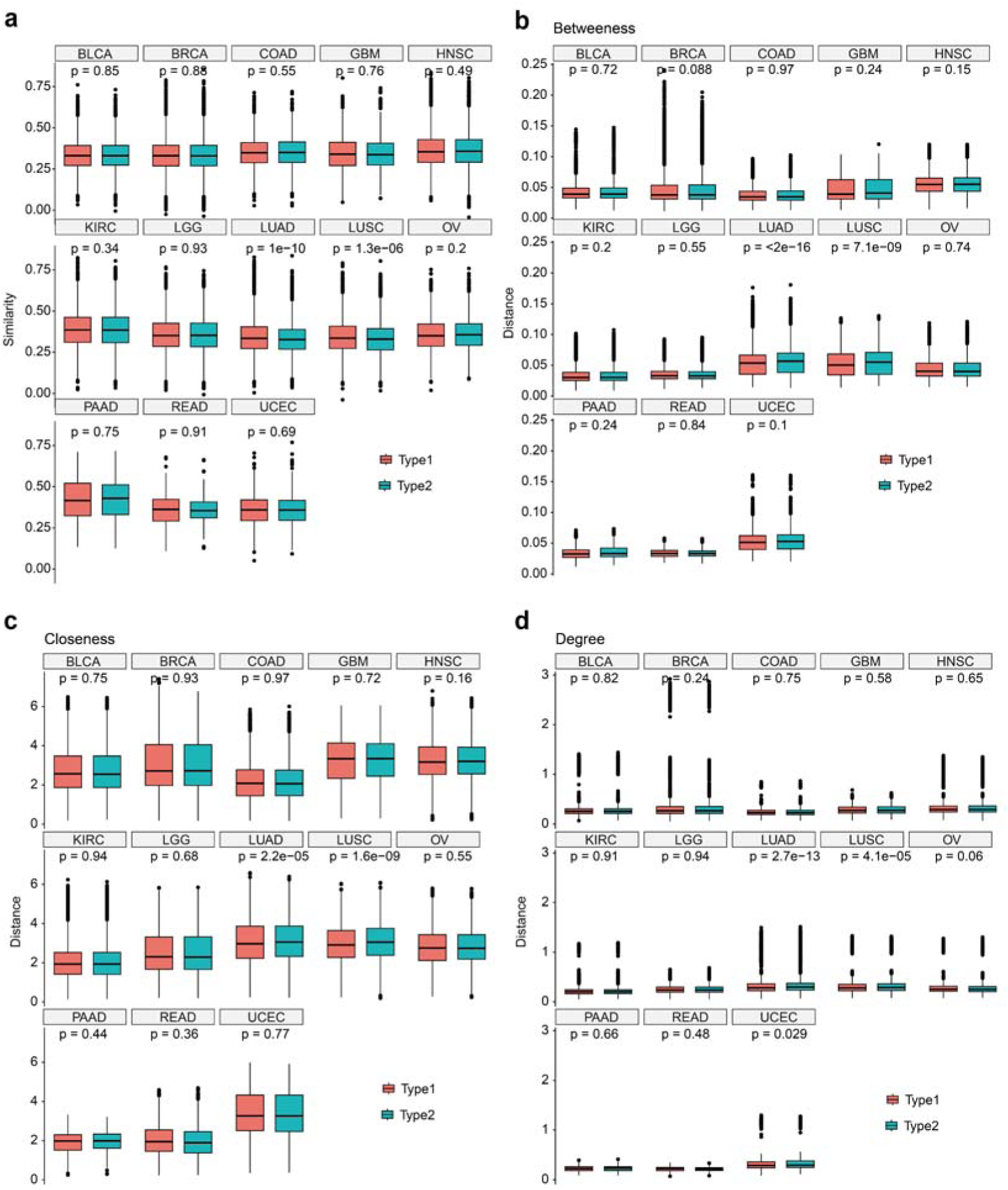
Comparison of enzyme network similarity and distance between samples with different hypoxic states. Type 1 were the similarity or distance of enzyme networks between samples in the same hypoxic state and Type 2 were the similarity or distance of enzyme networks between samples in different hypoxic states. **a,** Comparison of enzyme network similarity between samples with different hypoxic states in different cancer types. We used Graph2Vec, an algorithm designed to learn vector representations for whole graphs, to compute the embedding vector for each sample’s enzyme network. Then we use cosine similarity to calculate the similarity between pairs of enzyme networks. The gene-wise vector of centrality measures of each sample sepcific enzyme network were retrieved and computed the euclidean distance between these vectors of pairwise samples. Centrality measures included betweeness centrality **(b),** closeness centrality **(c)** and degree centrality **(d).** The sample sizes for different tumor types are detailed in **Supplementary Table 2.**

**Extended Data Fig. 2.**
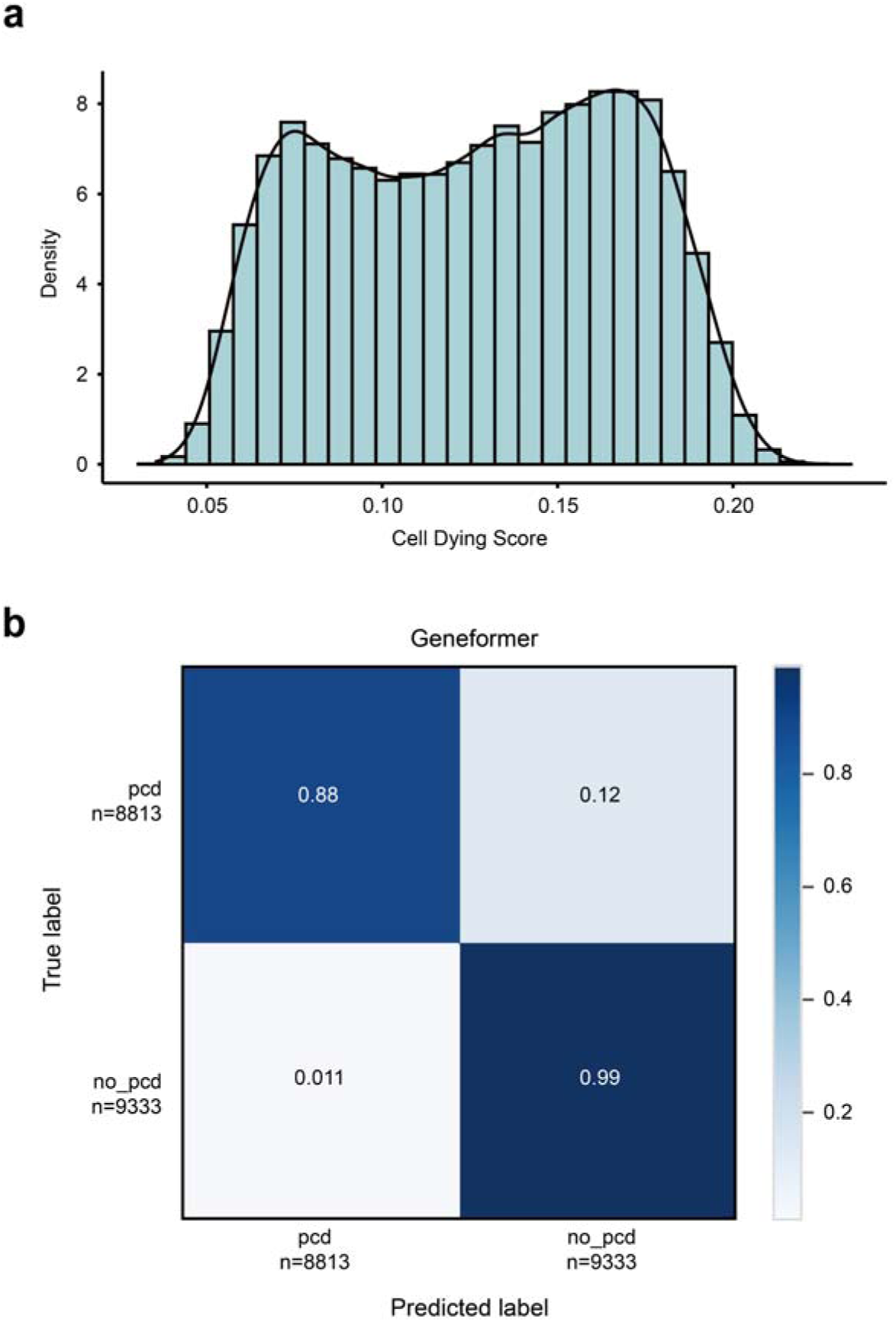
Performance of fine-tuned model. **a,** Distribution of cell dying score quantified by AUCell. **b,** Confusion matrix of fine-tuned model prediction on the independent test set (data obtained from Qian, J. et al).

**Extended Data Fig. 3.**
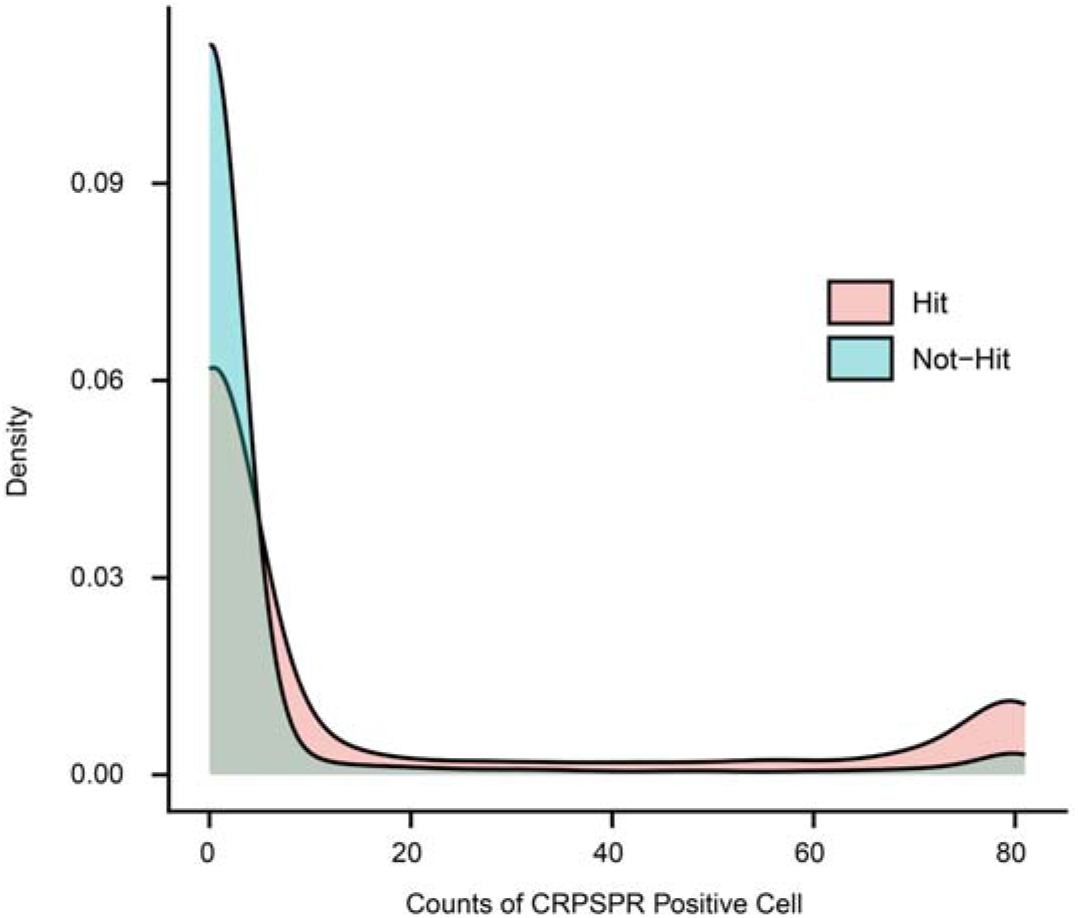
Comparison of the density distribution of CRISPR-positive cells between *in silico* perturbation hit and non-hit genes. *In silico* perturbation hit genes were defined as the effect value was > 0, and the FDR was < 0.05 (the effect value refers to the magnitude of cell embedding shift from the non-dying state to the dying state after deleting the specific gene). If a gene’s dependency score in a cell is >0.8, the cell is marked as a positive cell for that specific gene.

**Extended Data Fig. 4.**
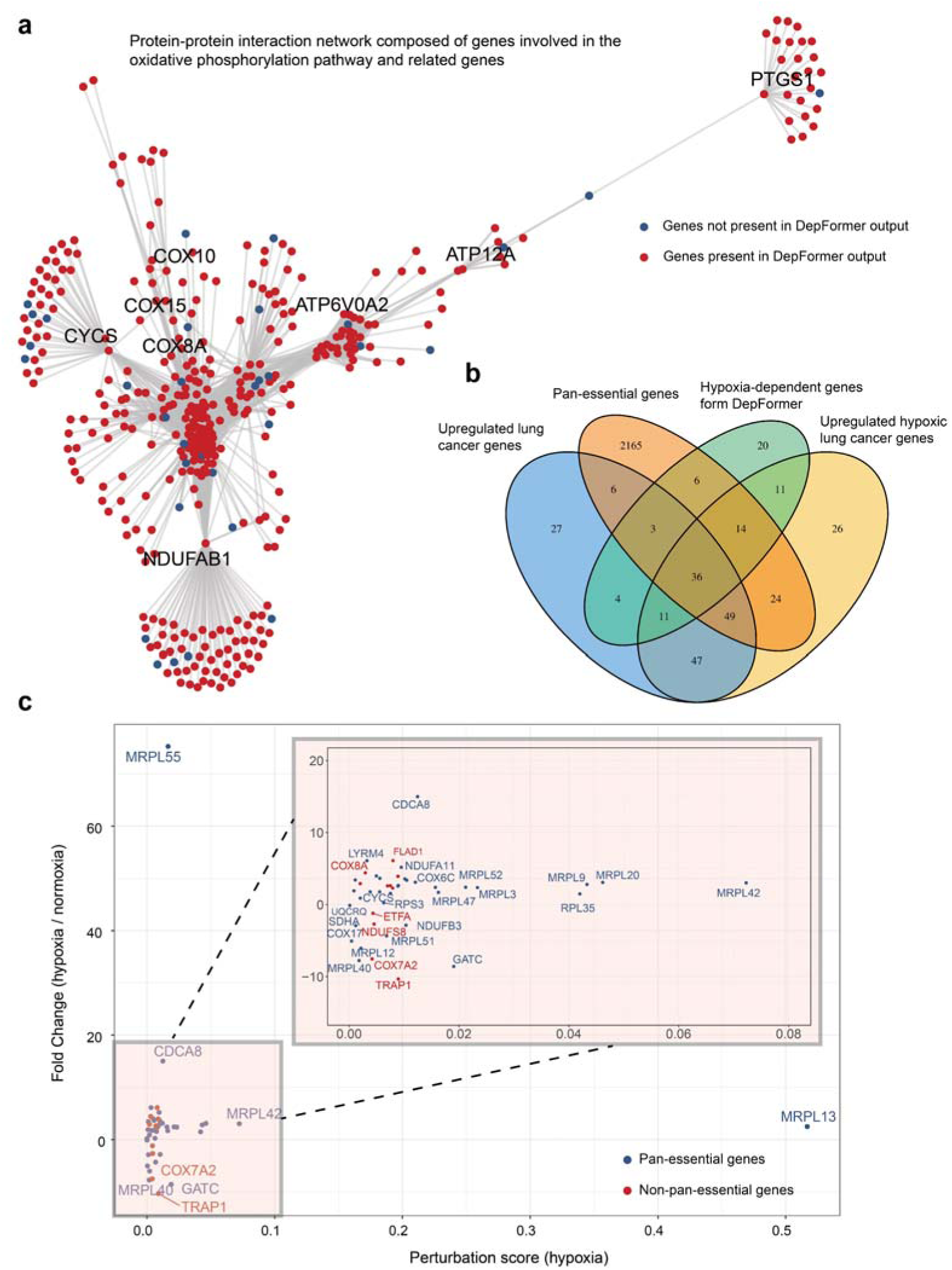
Genes specifically required for the survival of hypoxic tumor cells. **a,** A protein-protein interaction network consisting of oxidative phosphorylation pathway genes and genes that interact with these genes, in which genes identified by DepFormerare labeled in red. **b**, The Venn diagram illustrates the selection of key candidate genes associated with oxidative phosphorylation that play significant roles in tumor adaptation to hypoxia. A total of 11 OXPHOS-related genes were identified from the output of DepFormer based on the following criteria: 1) genes that are predicted by DepFormer as hypoxic lung cancer cell dependent; 2) genes that are significantly up-regulated in lung cancer compared to normal lung tissues; 3) genes that are significantly up-regulated in hypoxic lung cancer relative to non-hypoxic lung cancer; and 4) genes that are not classified as pan-essential genes for cellular viability. The perturbation scores for these genes are illustrated in **Fig. 1g. c,** A total of 47 genes that meet the first three criteria outlined in **Extended Data Fig. 4b** (excluding pan-essential genes) were identified. These genes are significantly up-regulated in tumors, particularly in hypoxic tumors, and have been predicted by DepFormer to be involved in hypoxia adaptation. Among these, FLAD1 stands out as a non-pan-essential gene with a high perturbation score in hypoxic cells, but not in non-hypoxic cells. Red dots represent non-pan-essential genes, and blue dots indicate pan-essential genes.

**Extended Data Fig. 5.**
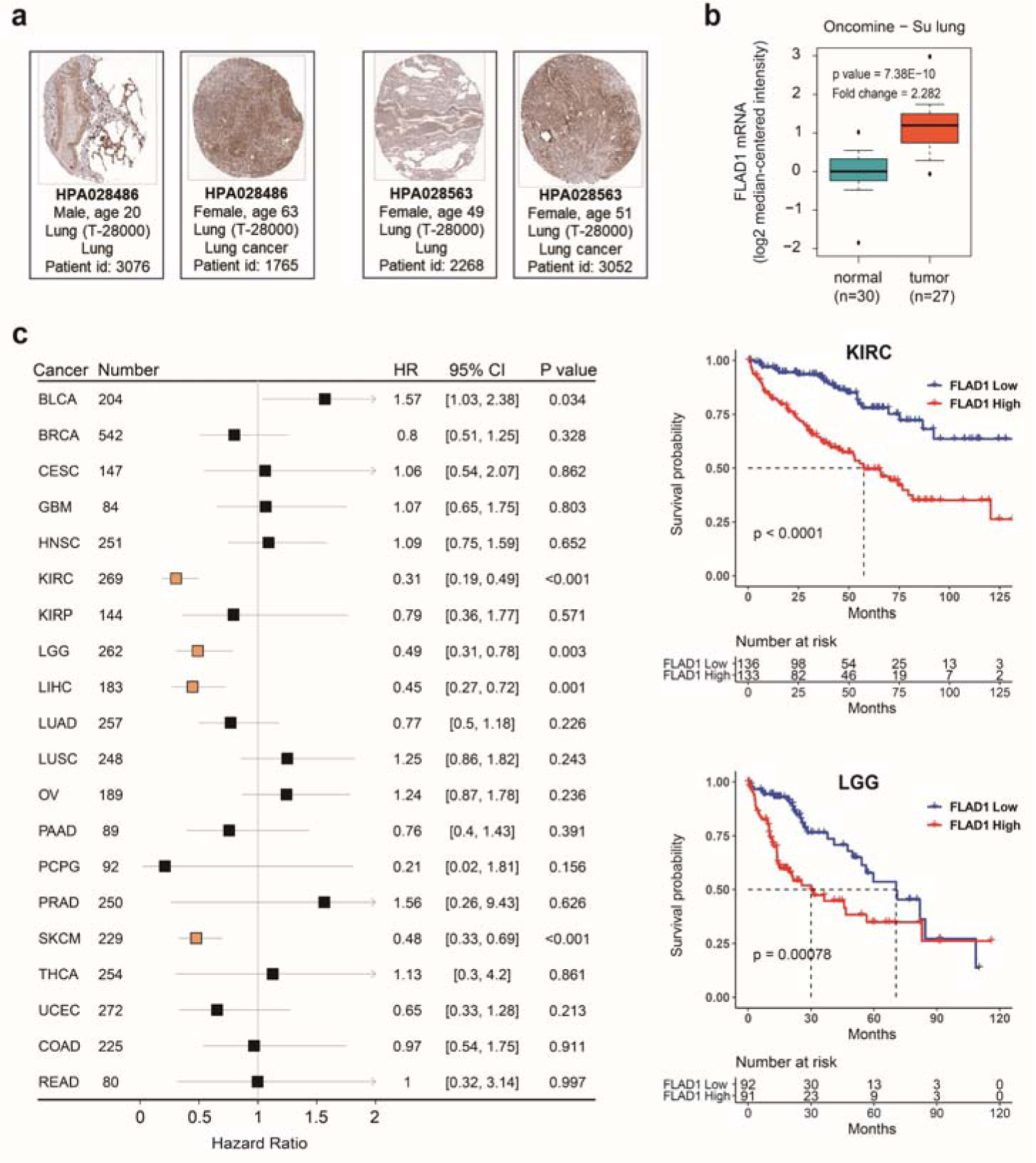
Protein expression characteristics of FLAD1 and its association with tumor prognosis. **a**, Representative immunohistochemical (IHC) images of FLAD1 in lung cancer and normal tissue obtained from the Human Protein Atlas (HPA) database. Quantification of positive cells was performed using ImageJ. **b**, Comparison of FLAD1 mRNA levels between lung adenocarcinoma and corresponding normal tissues in an independent dataset (Su Lung Statistics) from the Oncomine database. **c**, The forest plot on the left illustrates the hazard ratios for various tumors within the TCGA database. Kaplan-Meier survival curves for overall survival rates among KIRC, and LGG patients with high versus low FLAD1 expression in the TCGA database.

**Extended Data Fig. 6.**
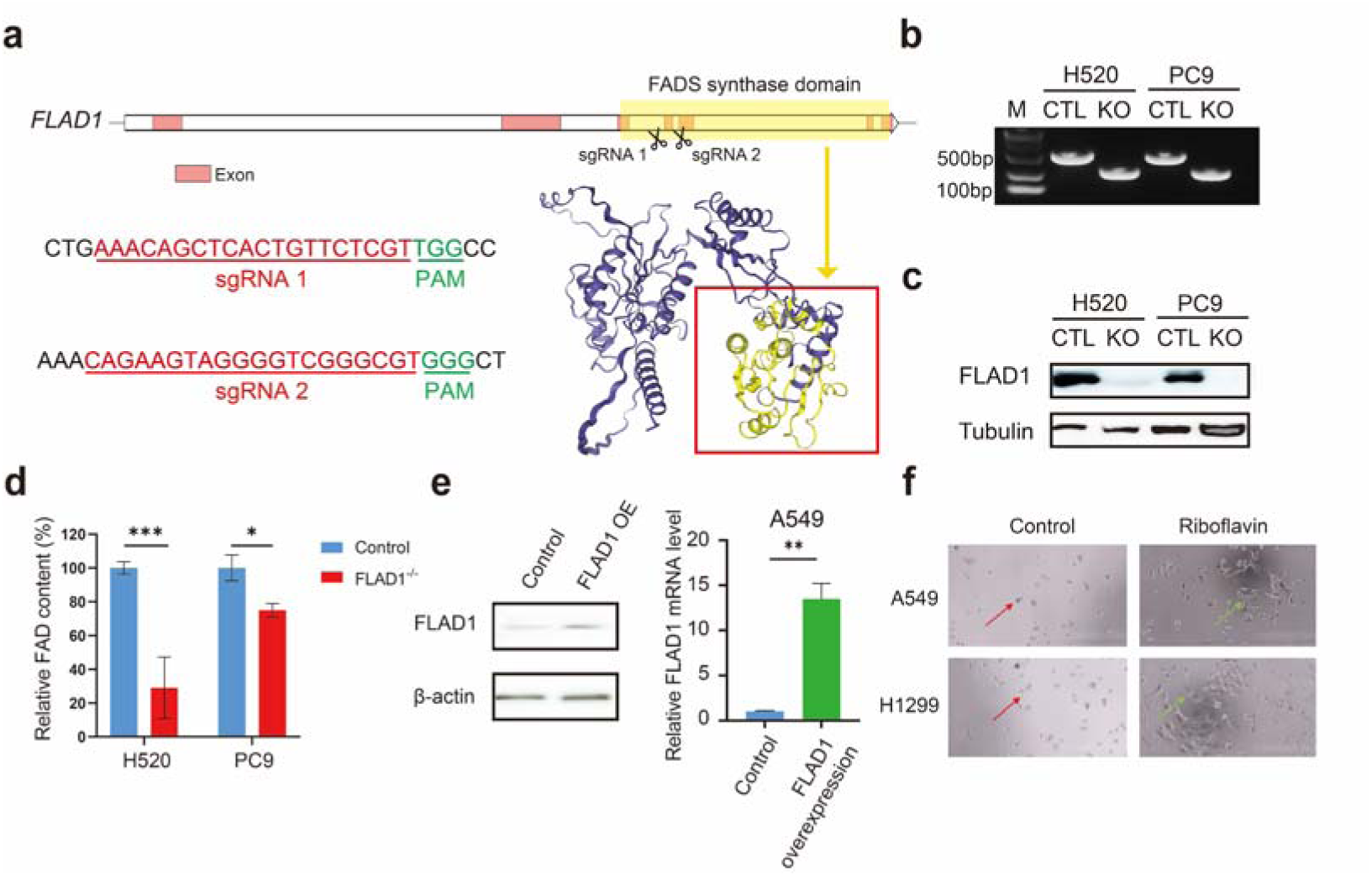
Construction and validation of FLAD1 knockout and overexpressing cell lines. **a**, Schematic representation of the CRISPR targeting the FAD synthesis functional domain (yellow region) of *FLAD1*. FLAD1 knockout was achieved using two sgRNAs, indicated by red bases, with the PAM sequences highlighted in green. **b**, PCR verification of *FLAD1* knockout cell lines. **c**, Western blot analysis for the confirmation of FLAD1 knockout. CTL: control cells; KO: knockout cells. **d**, Relative intracellular FAD concentrations (% to control) of control and *FLAD1* knockout cells. **e**, Western blot and qPCR analysis of FLAD1 protein and mRNA levels in A549 cells overexpressing *FLAD1* via lentivirus. OE: *FLAD1* overexpression. **f**, Representative images showing the proliferation and growth status of A549 and H1299 under hypoxia following 0.4 uM riboflavin supplementation. The red arrow indicates the growth status of cells under 1% oxygen conditions without the addition of riboflavin, whereas the green arrow signifies the growth status of cells with riboflavin supplementation under the same hypoxic conditions.

**Extended Data Fig. 7.**
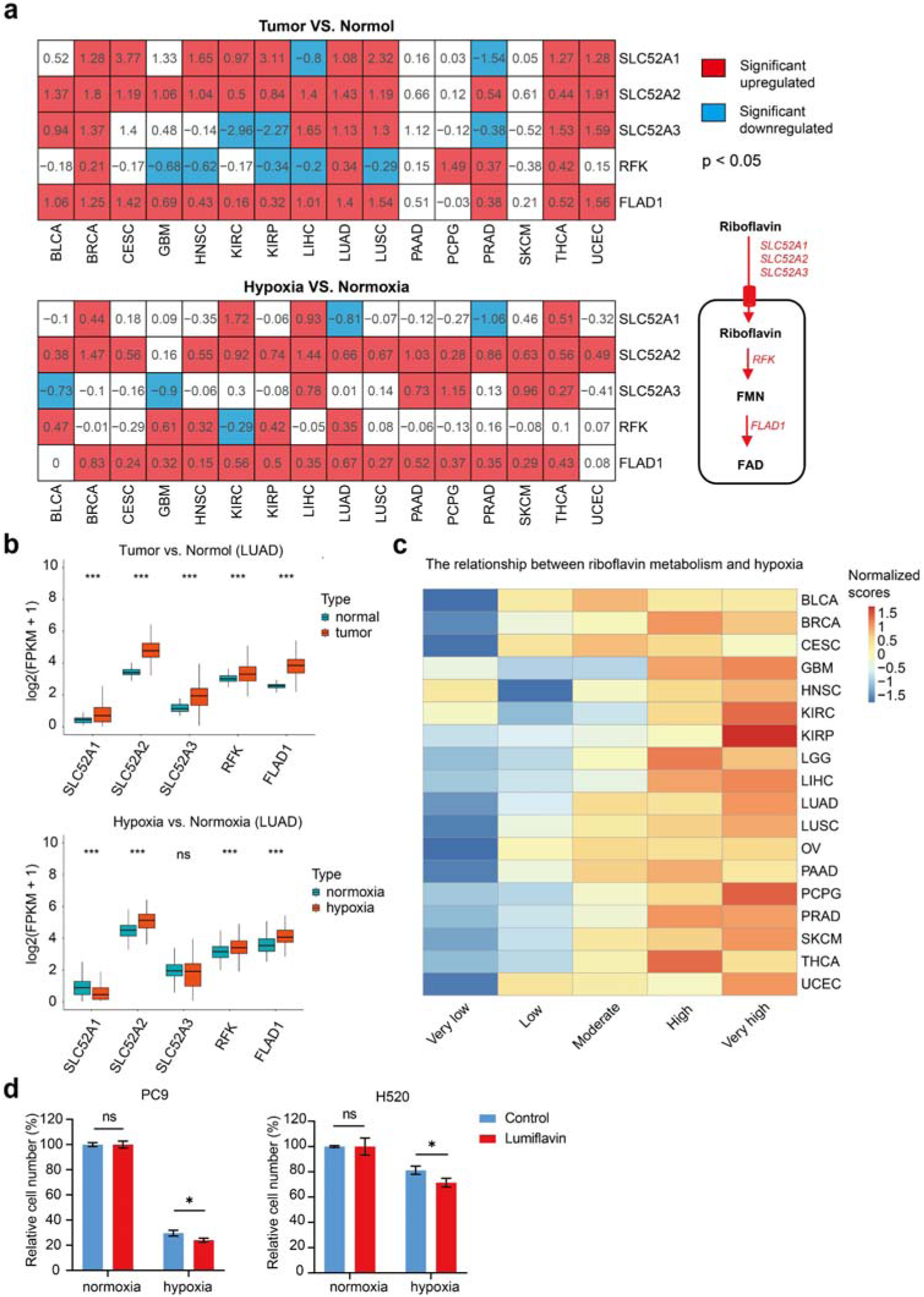
Aberrant activation of riboflavin metabolism in hypoxic tumor samples. **a**, Expression of riboflavin metabolism-related genes in tumors compared to normal tissues (top), and in hypoxic tumors versus non-hypoxic tumors (bottom) as per the TCGA database. Significance is indicated by p<0.05, and the numbers on the squares represent the log2(Fold Change). **b**, Differential expression of riboflavin metabolism-related genes in lung adenocarcinoma (LUAD). The upper panel displays the expression patterns of genes involved in riboflavin metabolism in samples from TCGA database, contrasting LUAD tissues with normal counterparts. The lower panel contrasts the expression profiles of these genes between hypoxic and non-hypoxic LUAD samples. Data are presented as median values (Q1, Q3). **c**, Riboflavin metabolism levels across tumor samples with varying degrees of hypoxia. Tumor samples of each type are stratified into quintiles based on the Buffa hypoxia score. These quintiles are then labeled according to the degree of hypoxia as “Very low,” “Low,” “Moderate,” “High,” and “Very high.” **d**, Effect of lumiflavin on the proliferation of H520 (50 uM lumiflavin) and PC9 (10 uM lumiflavin) cells under normoxic and hypoxic conditions.

**Extended Data Fig. 8.**
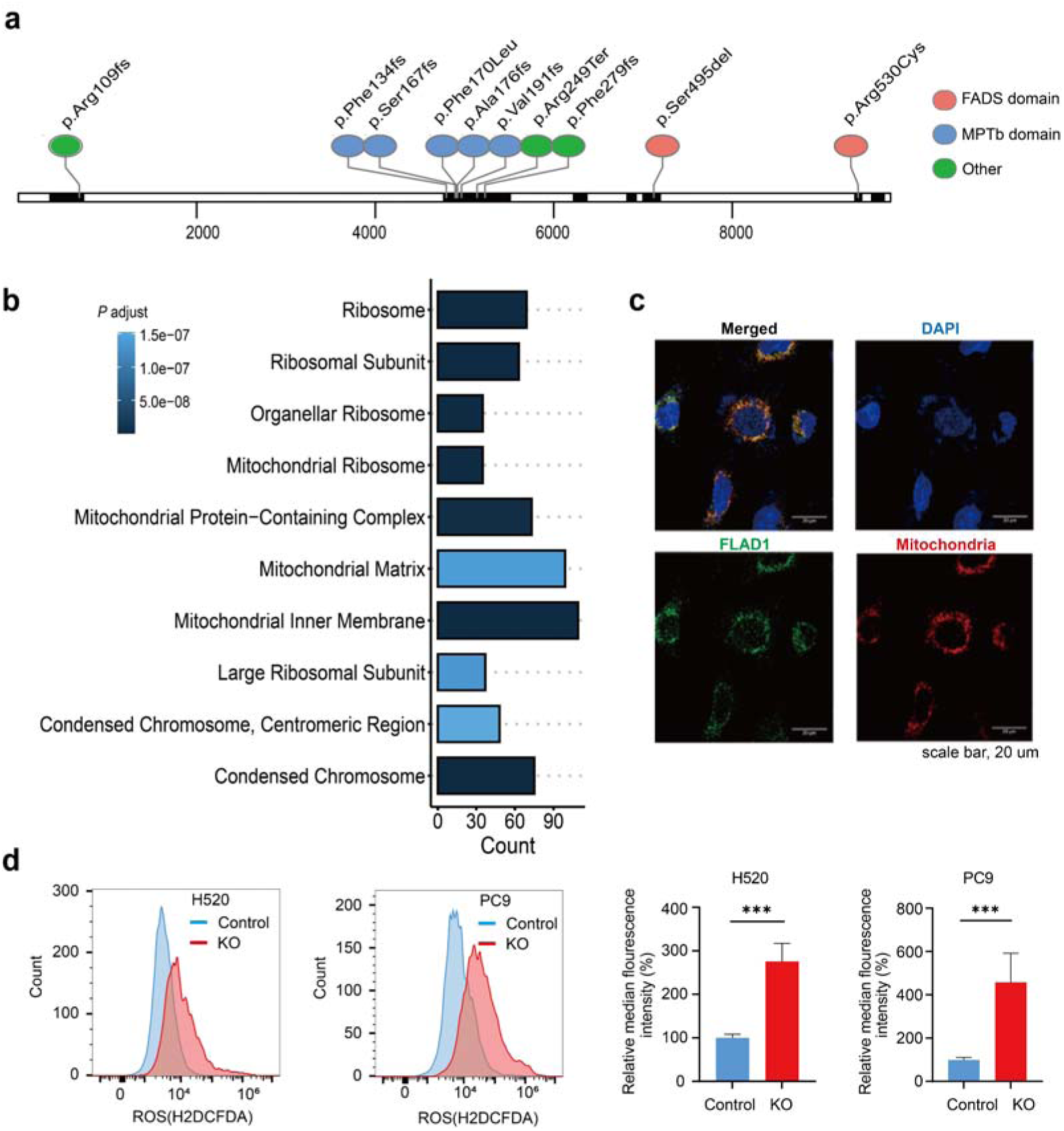
FLAD1 is associated with mitochondrial function. **a,** The mutation sites of FLAD1 in the human genetic disorder MADD are charted based on data from the Malacards database. Red markers denote mutations located within the FAD synthesis (FADS) domain of FLAD1, blue markers highlight mutations within the molybdopterin-binding (MPTb) domain, and green markers indicate mutations at other sites within the protein. The delineation of different functional domains is determined according to the UniProt database. **b**, GO analysis results of upregulated genes in lung cancer patients with high FLAD1 expression in the TCGA database. The figure illustrates the cellular localization of upregulated genes, highlighting the top ten most significantly enriched results, as determined by adjusted p-values. These results underscore the prominent subcellular compartments or pathways where the upregulated genes are predominantly involved. The horizontal axis denotes the number of genes enriched. **c**, Localization of FLAD1 in tumor cell line A549. Green indicates FLAD1 protein, red indicates mitochondria, and blue indicates cell nuclei stained with DAPI. Scale bar, 20 μm. **d**, The difference in ROS levels between control and *FLAD1* knockout (KO) tumor cells under normoxia. The histogram on the left displays the levels of ROS in control and KO groups, with blue denoting the control group and red representing the KO group. The bar graph on the right indicates the relative median fluorescence intensity in H520 and PC9 cells, comparing the KO group to the respective control group, thereby quantifying the degree of change in ROS levels attributable to the knockout condition.

**Extended Data Fig. 9.**
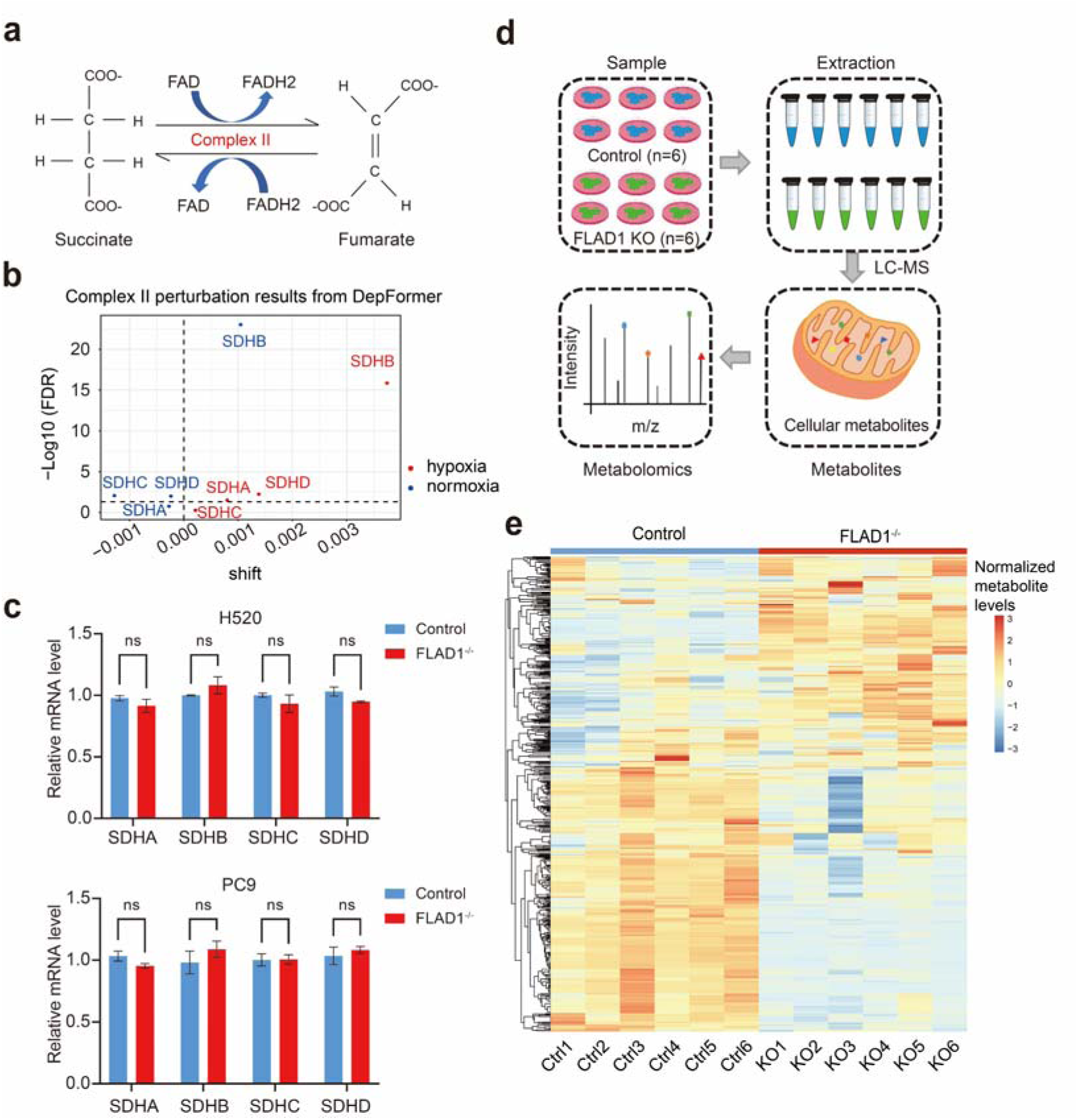
FLAD1 facilitates cellular adaptation to hypoxia via mitochondria complex II. **a**, A schematic illustrating the redox state changes of FAD on complex II under two different activity (succinate dehydrogenase and fumarate reductase) conditions. Under the condition of succinate dehydrogenase activity, FAD is reduced from its oxidation state(FAD) to FADH2, after which FADH2 can transfer electrons to the electron transfer chain to generate energy. Under the condition of fumarate reductase activity, FADH2 is oxidized to FAD, accepting electrons to promote the reduction reaction of fumarate. **b**, The *in silico* perturbation results for key genes of mitochondrial complex II outputted by DepFormer are presented. The shift value refers to the magnitude of cell embedding shift from the non-dying state to the dying state after deleting the specific gene. Red dots represent perturbation results for hypoxic lung cancer cells, while blue dots indicate results for non-hypoxic lung cancer cells. **c**, qPCR analysis of mRNA levels of the four main subunits of complex II in H520 and PC9 cells following *FLAD1* knockout. **d**, Schematic representation of the untargeted metabolomics workflow. **e**, Clustering results of metabolite expression levels between *FLAD1* wild-type control and knockout groups under hypoxia.

**Extended Data Fig. 10.**
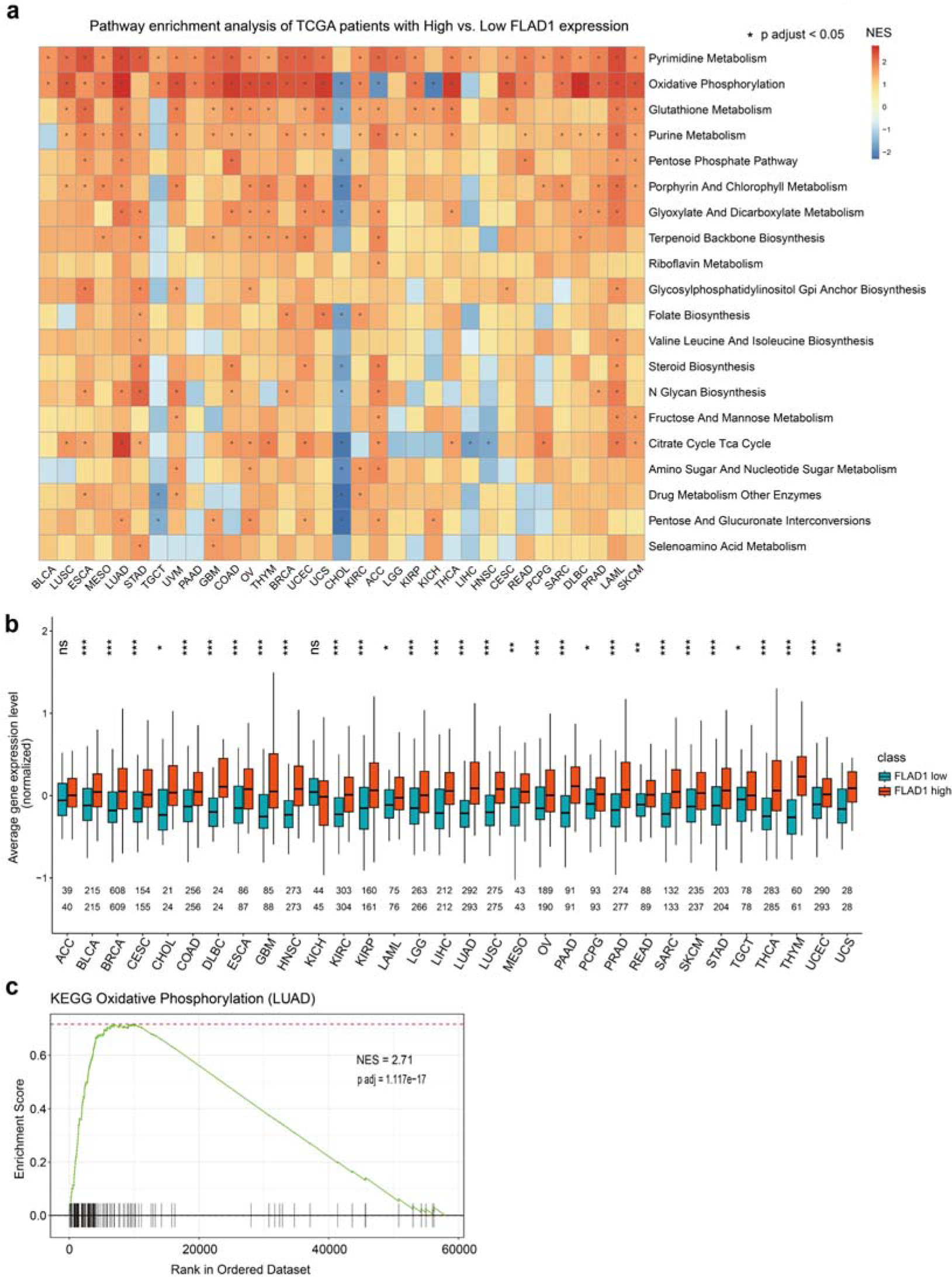
FLAD1 expression levels correlate with multiple cellular metabolic pathways in TCGA patients. **a**, GSEA of up-regulated genes in the high FLAD1 expression group across various tumor types in TCGA. Asterisks indicate an adjusted p-value of less than 0.05. The color blocks represent normalized enrichment score. **b**, Average expression levels of oxidative phosphorylation genes in high vs. low FLAD1 expression groups across multiple tumor types. The two rows of data above the axis represent the number of samples in the FLAD1-high and FLAD1-low groups, respectively. **c**, GSEA demonstrating enrichment of oxidative phosphorylation gene sets in lung adenocarcinoma patients with high FLAD1 expression.

**Extended Data Fig. 11.**
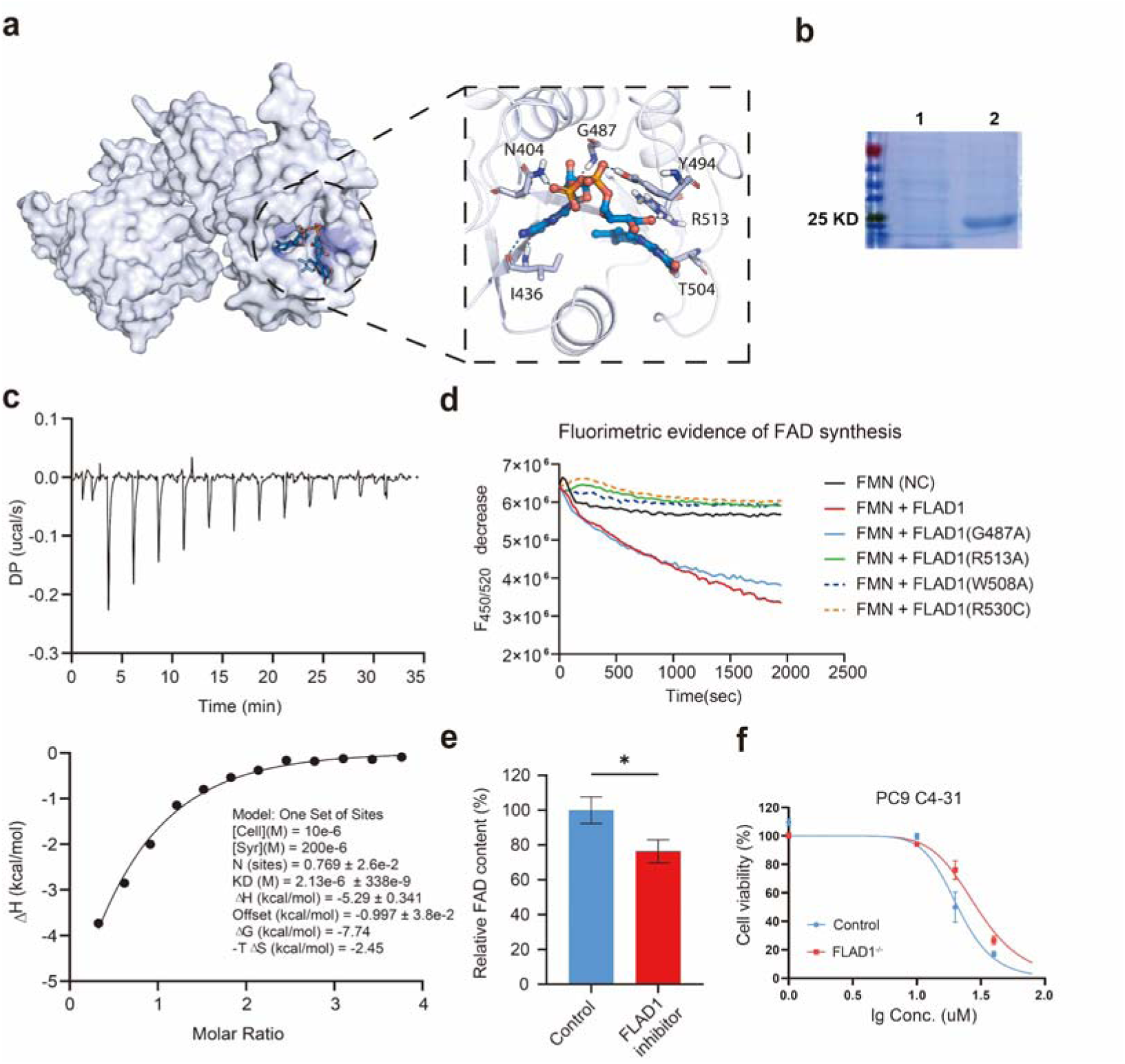
In vitro selection of FLAD1 inhibitors. **a**, Binding mode of FLAD1 (light blue) and FAD (marine). FLAD1 was shown in surface mode in the left subplot. Active residues were shown as light blue sticks. **b,** Display of purified recombinant FLAD1 protein. The first lane of the protein gel indicates bacteria not induced with IPTG, while the second lane shows protein induced with IPTG and subsequently purified. **c**, ITC binding data of FMN titrated into recombinant FLAD1 protein. **d,** Establishment of a screening assay for FLAD1 inhibitors. Active FLAD1 catalyzes the conversion of highly FMN to the less fluorescent FAD. The activity of the in vitro purified recombinant FLAD1 protein is monitored by detecting the emitted light at 520 nm upon excitation at 450 nm, capitalizing on the differential fluorescence properties of FMN and FAD. **e,** Changes in intracellular FAD content after adding compound C4-31. **f,** IC50 of compound C4-31 on cell lines with wild-type FLAD1 and FLAD1 knockout.

Supplementary Table 1. Abbreviations for tumor types and the number of tumor samples from TCGA mRNA expression datasets.

Supplementary Table 2. The number of samples from different tumor types used in the enzyme network analysis.

Supplementary Table 3. Top 50 filtered templates for FLAD1 sequence.

Supplementary Table 4. Computational evaluation for the bindings between FLAD1 and compounds selected from Specs compound library.

Supplementary Table 5. Compounds modified at R1 position.

Supplementary Table 6. Compounds modified at R2 position.

Supplementary Table 7. Compounds modified at R1 and R2 position.

Supplementary Table 8. Compounds containing carboxyl hydrazide group screened in substructure search.

Supplementary Table 9. Alanine scanning result of complex of FLAD1 and C4-31.

Supplementary Table 10. ADMET prediction.

Supplementary Table 11. Primers for qPCR.

## Notes

### Competing Interest Statement

The authors have declared no competing interest.

