## Supplemental Table for "Tumor hypoxia adaptation depends on FLAD1 mediated mitochondrial metabolic reprogramming"

**Supplementary Table 1. Abbreviations for tumor types and the number of tumor samples from TCGA mRNA expression datasets.**

| Study Abbreviation | Study Name | Number of tumor samples | Number of non-tumor samples |
|---|---|---|---|
| BLCA | Bladder Urothelial Carcinoma | 411 | 19 |
| BRCA | Breast invasive carcinoma | 1104 | 113 |
| CESC | Cervical squamous cell carcinoma and | 306 | 3 |
| COAD | Colon adenocarcinoma | 471 | 41 |
| GBM | Glioblastoma multiforme | 168 | 5 |
| HNSC | Head and Neck squamous cell | 502 | 44 |
| KIRC | Kidney renal clear cell carcinoma | 535 | 72 |
| KIRP | Kidney renal papillary cell carcinoma | 289 | 32 |
| LGG | Brain Lower Grade Glioma | 529 | 0 |
| LIHC | Liver hepatocellular carcinoma | 374 | 50 |
| LUAD | Lung adenocarcinoma | 526 | 59 |
| LUSC | Lung squamous cell carcinoma | 501 | 49 |
| OV | Ovarian serous cystadenocarcinoma | 379 | 0 |
| PAAD | Pancreatic adenocarcinoma | 178 | 4 |
| PCPG | Pheochromocytoma and Paraganglioma | 183 | 3 |
| PRAD | Prostate adenocarcinoma | 499 | 52 |
| READ | Rectum adenocarcinoma | 167 | 10 |
| SKCM | Skin Cutaneous Melanoma | 471 | 1 |
| THCA | Thyroid carcinoma | 510 | 58 |
| UCEC | Uterine Corpus Endometrial Carcinoma | 548 | 35 |

**Supplementary Table 2. The number of samples from different tumor types used in the enzyme network analysis.**

| Tumor type | Type1 | Type2 |
|---|---|---|
| BLCA | 5790 | 5832 |
| BRCA | 60741 | 60984 |
| COAD | 2686 | 2756 |
| GBM | 1259 | 1295 |
| HNSC | 11447 | 11556 |
| KIRC | 10908 | 11016 |
| LGG | 6014 | 6075 |
| LUAD | 9119 | 9207 |
| LUSC | 5182 | 5254 |
| OV | 2555 | 2494 |
| PAAD | 306 | 324 |
| READ | 213 | 221 |
| UCEC | 849 | 858 |

**Supplementary Table 3. Top 50 filtered templates for FLAD1 sequence.**

| Template | Seq Identit | Oligo-state | QSQE | Found by | Method | Resolution | Seq Similarity | Coverage | Description |
|---|---|---|---|---|---|---|---|---|---|
| Q8NFF5.1.A | 100 | monomer | - | AFDB search | AlphaFold v2 | NA | 0.62 | 1 | FAD synthase |
| 3kbq.1.B | 37.93 | homo- | 0.19 | BLAST | X-ray | 2.00Å | 0.38 | 0.25 | Protein Ta0487 |
| 3kbq.1.A | 37.93 | homo- | 0.19 | BLAST | X-ray | 2.00Å | 0.38 | 0.25 | Protein Ta0487 |
| 2wsi.1.A | 37.31 | monomer | - | BLAST | X-ray | 1.90Å | 0.39 | 0.34 | FAD SYNTHETASE |
| 3kbq.1.A | 33.55 | homo- | 0.19 | HHblits | X-ray | 2.00Å | 0.35 | 0.26 | Protein Ta0487 |
| 3kbq.1.B | 33.55 | homo- | 0.19 | HHblits | X-ray | 2.00Å | 0.35 | 0.26 | Protein Ta0487 |
| 2wsi.1.A | 33.02 | monomer | - | HHblits | X-ray | 1.90Å | 0.37 | 0.37 | FAD SYNTHETASE |
| 3g6k.3.A | 31.73 | monomer | - | BLAST | X-ray | 1.35Å | 0.37 | 0.35 | FMN adenylyltransferase |
| 3g5a.6.A | 31.73 | monomer | - | BLAST | X-ray | 1.95Å | 0.37 | 0.35 | FMN adenylyltransferase |
| 3g5a.5.A | 31.73 | monomer | - | BLAST | X-ray | 1.95Å | 0.37 | 0.35 | FMN adenylyltransferase |
| 3g5a.2.A | 31.73 | monomer | - | BLAST | X-ray | 1.95Å | 0.37 | 0.35 | FMN adenylyltransferase |
| 3g6k.3.A | 31.63 | monomer | - | HHblits | X-ray | 1.35Å | 0.36 | 0.37 | FMN adenylyltransferase |
| 3g5a.6.A | 31.63 | monomer | - | HHblits | X-ray | 1.95Å | 0.36 | 0.37 | FMN adenylyltransferase |
| 3g5a.5.A | 31.63 | monomer | - | HHblits | X-ray | 1.95Å | 0.36 | 0.37 | FMN adenylyltransferase |
| 3g5a.2.A | 31.63 | monomer | - | HHblits | X-ray | 1.95Å | 0.36 | 0.37 | FMN adenylyltransferase |
| 4kkv.1.A | 31.25 | monomer | - | BLAST | X-ray | 1.74Å | 0.37 | 0.35 | Similar to uniprot\|P38913 Saccharomyces cerevisiae YDL045c FAD synthetase |
| 4kkv.1.A | 31.16 | monomer | - | HHblits | X-ray | 1.74Å | 0.36 | 0.37 | Similar to uniprot\|P38913 Saccharomyces cerevisiae YDL045c FAD synthetase |
| 4ct9.1.A | 29.39 | homo- | 0.18 | HHblits | X-ray | 2.14Å | 0.33 | 0.39 | CINA-LIKE PROTEIN |
| 4uux.1.A | 29.39 | homo- | 0.16 | HHblits | X-ray | 1.99Å | 0.33 | 0.39 | CINA |
| 4cta.1.A | 29.39 | monomer | - | HHblits | X-ray | 2.21Å | 0.32 | 0.39 | CINA-LIKE PROTEIN |
| 4cta.1.B | 29.39 | monomer | - | HHblits | X-ray | 2.21Å | 0.33 | 0.39 | CINA-LIKE PROTEIN |
| 4uux.1.B | 29.39 | homo- | 0.16 | HHblits | X-ray | 1.99Å | 0.33 | 0.39 | CINA |
| 4bwv.1.A | 24.88 | homo-dimer | - | BLAST | X-ray | 1.80Å | 0.34 | 0.36 | PHOSPHOADENOSINE-PHOSPHOSULPHATE |
| 4bwv.1.A | 22.6 | homo-dimer | - | HHblits | X-ray | 1.80Å | 0.32 | 0.3 | PHOSPHOADENOSINE-PHOSPHOSULPHATE |
| 1di6.1.A | 21.91 | homo-trimer | - | HHblits | X-ray | 1.45Å | 0.3 | 0.3 | MOLYBDENUM COFACTOR BIOSYNTHETIC ENZYME |
| 1zun.1.A | 21.46 | monomer | - | HHblits | X-ray | 2.70Å | 0.31 | 0.35 | Sulfate adenylyltransferase subunit 2 |

| Template | Seq Identit | Oligo-state | QSQE | Found by | Method | Resolution | Seq Similarity | Coverage | Description |
|---|---|---|---|---|---|---|---|---|---|
| 6mr3.2.A | 20 | monomer | - | HHblits | X-ray | 2.05Å | 0.3 | 0.38 | Putative competence-damage inducible protein |
| 6mr3.1.B | 20 | monomer | - | HHblits | X-ray | 2.05Å | 0.3 | 0.38 | Putative competence-damage inducible protein |
| 6mr3.1.A | 20 | monomer | - | HHblits | X-ray | 2.05Å | 0.3 | 0.38 | Putative competence-damage inducible protein |
| 2oq2.1.A | 19.19 | homo-dimer | - | HHblits | X-ray | 2.10Å | 0.3 | 0.29 | Phosphoadenosine phosphosulfate reductase |
| 2goy.1.B | 19.05 | homo-tetramer | - | HHblits | X-ray | 2.70Å | 0.31 | 0.29 | adenosine phosphosulfate reductase |
| 2goy.1.A | 19.05 | homo-tetramer | - | HHblits | X-ray | 2.70Å | 0.31 | 0.29 | adenosine phosphosulfate reductase |
| 1gpm.1.D | 19.02 | monomer | - | HHblits | X-ray | 2.20Å | 0.29 | 0.28 | GMP SYNTHETASE |
| 1gpm.1.A | 19.02 | monomer | - | HHblits | X-ray | 2.20Å | 0.29 | 0.28 | GMP SYNTHETASE |
| 1gpm.1.C | 19.02 | monomer | - | HHblits | X-ray | 2.20Å | 0.29 | 0.28 | GMP SYNTHETASE |
| 6vpu.2.A | 18.93 | monomer | - | HHblits | X-ray | 1.90Å | 0.31 | 0.29 | Phosphoadenosine phosphosulfate reductase |
| 6vpu.5.A | 18.93 | monomer | - | HHblits | X-ray | 1.90Å | 0.31 | 0.29 | Phosphoadenosine phosphosulfate reductase |
| 6vpu.1.A | 18.93 | monomer | - | HHblits | X-ray | 1.90Å | 0.31 | 0.29 | Phosphoadenosine phosphosulfate reductase |
| 6vpu.8.A | 18.93 | monomer | - | HHblits | X-ray | 1.90Å | 0.31 | 0.29 | Phosphoadenosine phosphosulfate reductase |
| 6vpu.6.A | 18.93 | monomer | - | HHblits | X-ray | 1.90Å | 0.31 | 0.29 | Phosphoadenosine phosphosulfate reductase |
| 7rge.1.A | 18.9 | monomer | - | HHblits | X-ray | 2.38Å | 0.31 | 0.28 | 3'-phosphoadenylylsulfate reductase |
| 2o8v.1.A | 18.4 | homo-dimer | - | HHblits | X-ray | 3.00Å | 0.3 | 0.28 | Phosphoadenosine phosphosulfate reductase |
| 1sur.1.A | 18.35 | homo- | - | HHblits | X-ray | 2.00Å | 0.31 | 0.27 | PAPS REDUCTASE |
| 7sbc.1.A | 16.77 | homo- | - | HHblits | X-ray | 1.95Å | 0.3 | 0.27 | GMP synthase [glutamine- |
| 3a4i.1.B | 16.56 | monomer | - | HHblits | X-ray | 1.79Å | 0.28 | 0.28 | GMP synthase [glutamine-hydrolyzing] subunit B |
| 3a4i.1.A | 16.56 | monomer | - | HHblits | X-ray | 1.79Å | 0.28 | 0.28 | GMP synthase [glutamine-hydrolyzing] subunit B |
| 6jp9.1.A | 15.85 | monomer | - | HHblits | X-ray | 2.10Å | 0.29 | 0.28 | GMP synthase [glutamine-hydrolyzing] subunit B |

| Template | Seq Identit | Oligo-state | QSQE | Found by | Method | Resolution | Seq Similarity | Coverage | Description |
|---|---|---|---|---|---|---|---|---|---|
| 6jp9.1.B | 15.85 | monomer | - | HHblits | X-ray | 2.10Å | 0.29 | 0.28 | GMP synthase [glutamine-hydrolyzing] subunit B |
| 7mo6.1.A | 13.86 | monomer | - | HHblits | X-ray | 2.30Å | 0.27 | 0.28 | GMP synthase [glutamine- |
| 7mo6.1.B | 13.86 | monomer | - | HHblits | X-ray | 2.30Å | 0.27 | 0.28 | GMP synthase [glutamine- |

**Supplementary Table 4. computational evaluation for the binding between FLAD1 and compounds selected from Specs compound library.**

| Compound ID | Compound Structure | Docking score | MM/GBSA ($\Delta G_{Bind}$) |
|---|---|---|---|
| FAD | 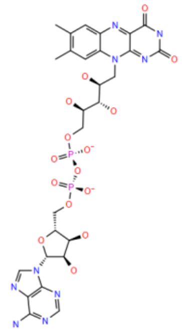 | -13.602 | -79.69 |
| C005 | 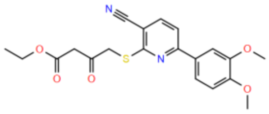 | -7.508 | -87.84 |
| C002 | 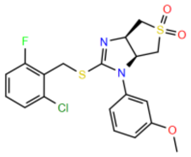 | -6.727 | -80.73 |
| C003 | 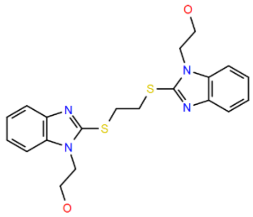 | -7.601 | -80.69 |
| C001 | 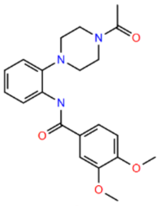 | -7.481 | -80.48 |
| C004 | 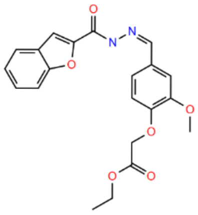 | -6.701 | -80.47 |
| C006 | 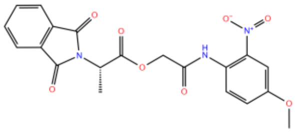 | -7.378 | -79.61 |

**Supplementary Table 5. Compounds modified at R1 position.**

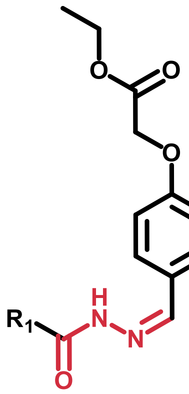


| Compound ID | Compound Structure | Docking score | MM/GBSA($\Delta G_{Bind}$) |
|---|---|---|---|
| C4-20 | 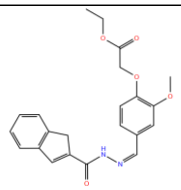 | -5.946 | -62.09 |
| C4-21 | 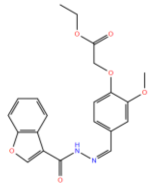 | -6 | -61.72 |
| C4-23 | 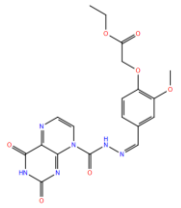 | -4.927 | -75.26 |
| C4-24 | 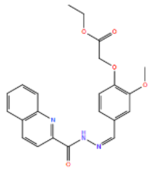 | -6.413 | -55.31 |
| C4-25 | 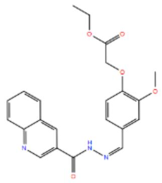 | -6.923 | -75.49 |
| C4-26 | 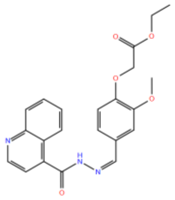 | -5.875 | -61.15 |
| C4-27 | 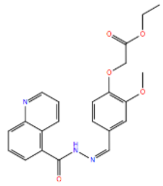 | -5.783 | -60.32 |

| | | | |
|---|---|---|---|
| C4-28 | 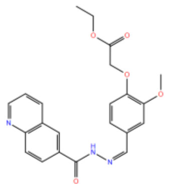 | -7.294 | -79.38 |
| C4-29 | 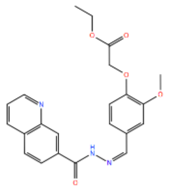 | -6.73 | -77.39 |
| C4-30 | 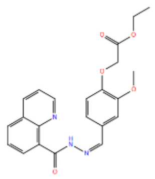 | -6.215 | -67.99 |
| C4-31 | 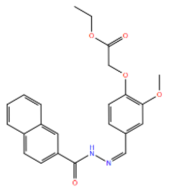 | -7.069 | -77.73 |
| C4-32 | 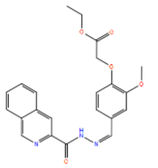 | -7.032 | -81.46 |
| C4-33 | 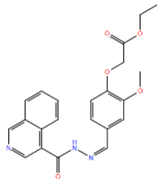 | -6.075 | -68.86 |
| C4-34 | 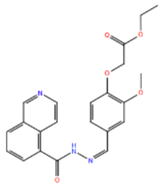 | -7.307 | -67.58 |
| C4-35 | 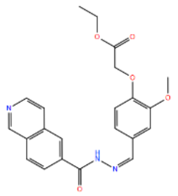 | -6.732 | -78.25 |
| C4-36 | 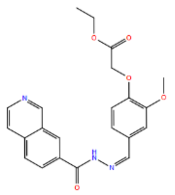 | -6.769 | -82.46 |

| | | | |
|---|---|---|---|
| C4-37 | 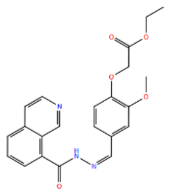 | -5.203 | -57.12 |

**Supplementary Table 6. Compounds modified at $R_2$ position.**

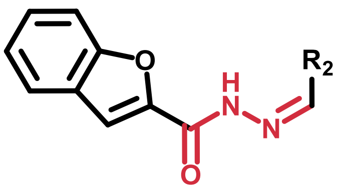


| Compound ID | Compound Structure | Docking score | MM/GBSA($\Delta G_{Bind}$) |
|---|---|---|---|
| C4-1 | 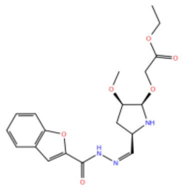 | -6.624 | -79.88 |
| C4-2 | 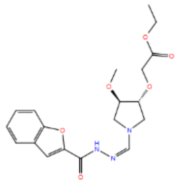 | -6.939 | -82.18 |
| C4-3 | 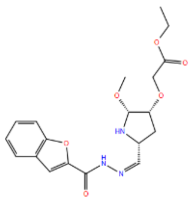 | -4.292 | -51.79 |
| C4-4 | 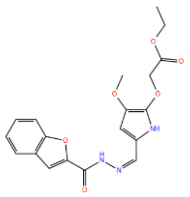 | -6.954 | -75.51 |
| C4-5 | 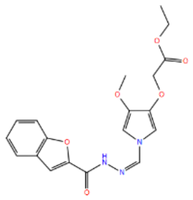 | -6.892 | -74.55 |
| C4-6 | 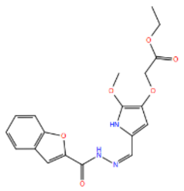 | -5.721 | -64.39 |
| C4-7 | 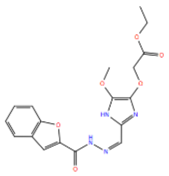 | -7.015 | -77.54 |

| | | | |
|---|---|---|---|
| C4-8 | 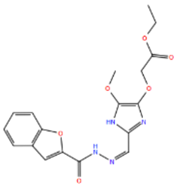 | -6.599 | -69.19 |
| C4-9 | 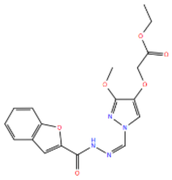 | -6.881 | -76.62 |
| C4-10 | 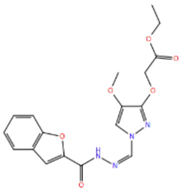 | -6.669 | -71.84 |
| C4-11 | 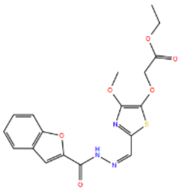 | -6.707 | -76.19 |
| C4-12 | 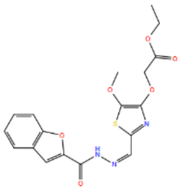 | -5.887 | -69.28 |
| C4-13 | 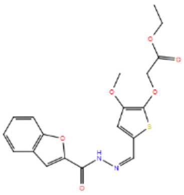 | -6.265 | -70.05 |
| C4-14 | 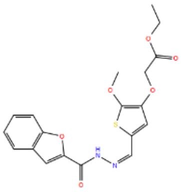 | -6.483 | -77.25 |
| C4-15 | 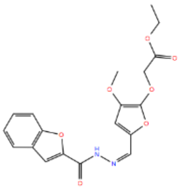 | -5.911 | -58.07 |
| C4-16 | 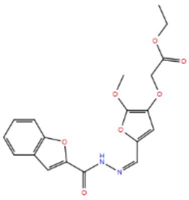 | -6.624 | -80.71 |

| | | | |
|---|---|---|---|
| C4-17 | 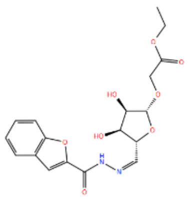 | -5.694 | -51.72 |
| C4-18 | 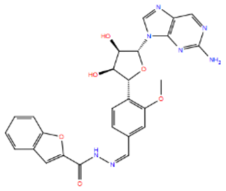 | -5.916 | -65.48 |
| C4-19 | 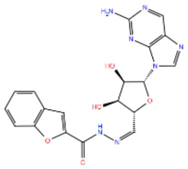 | -7.521 | -63.63 |
| C4-22 | 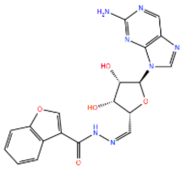 | -7.503 | -49.72 |

**Supplementary Table 7. Compounds modified at $R_1$ and $R_2$ position.**

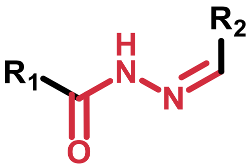


| Compound ID | Compound Structure | Docking score | MM/GBSA($\Delta G_{Bind}$) |
|---|---|---|---|
| C4-38 | 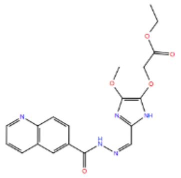 | -6.671 | -78.54 |

**Supplementary Table 8. Compounds containing carboxyl hydrazide group screened in substructure search.**

| Compound ID | Compound Structure | Docking score | MM/GBSA($\Delta G_{Bind}$) |
|---|---|---|---|
| C2-1 | 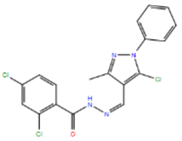 | -7.863 | -89 |
| C2-2 | 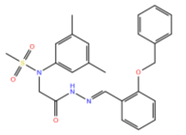 | -7.302 | -75.42 |
| C2-3 | 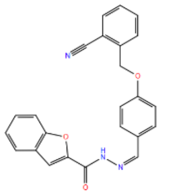 | -7.296 | -78.82 |
| C2-4 | 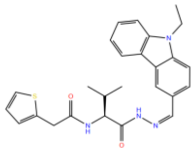 | -7.182 | -75.45 |
| C2-5 | 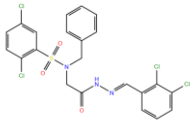 | -7.599 | -75.29 |
| C2-6 | 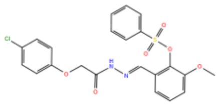 | -7.132 | -81.8 |
| C2-7 | 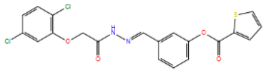 | -6.66 | -72.83 |
| C2-8 | 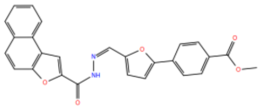 | -7.787 | -76.36 |
| C2-9 | 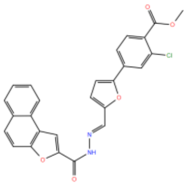 | -7.484 | -88.59 |
| C2-10 | 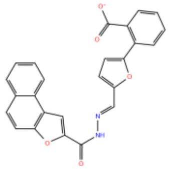 | -7.441 | -67.96 |

| | | | |
|---|---|---|---|
| C2-11 | 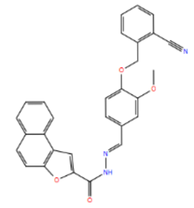 | -7.398 | -86.02 |
| C2-12 | 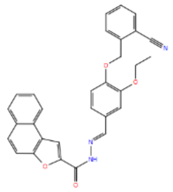 | -7.283 | -85.28 |
| C2-13 | 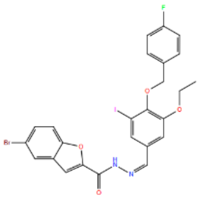 | -6.539 | -95.49 |
| C2-14 | 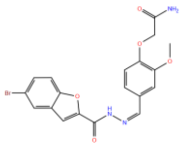 | -6.506 | -88.82 |
| C2-15 | 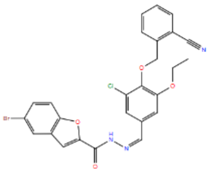 | -6.46 | -85.23 |

**Supplementary Table 9. Alanine scanning result of complex of FLAD1 and C4-31.**

| Residue | Original | Mutated | ∆Affinity | ∆Stability (solvated) |
|---|---|---|---|---|
| A:487 | GLY | ALA | 0.25 | 0.87 |
| A:409 | CYS | ALA | 1.25 | 1.71 |
| A:404 | ASN | ALA | 6.61 | 9.14 |
| A:508 | TRP | ALA | 10.67 | 14.7 |
| A:513 | ARG | ALA | 15.7 | 37.65 |

**Supplementary Table 10. ADMET prediction.**

| compound | C4-31 | Chicago Sky Blue |
|---|---|---|
| smiles | CCOC(=O)COc1ccc(/C=N\NC(=O)c2ccc3ccccc3c2)cc1OC | COC1=C(C=CC(=C1)C2=CC(=C(C=C2)N=NC3=C(C4=C(C=C3)C(=CC(=C4N)S(=O)(=O)[O-])S(=O)(=O)[O-])O)OC)N=NC5=C(C6=C(C=C5)C(=CC(=C6N)S(=O)(=O)[O-])S(=O)(=O)[O-])O.[Na+].[Na+].[Na+].[Na+] |
| LogS | -5.316 | -5.395 |
| LogD | 2.924 | 0.088 |
| LogP | 3.967 | 3.587 |
| Pgp-inh | 0.995 | 0.518 |
| Pgp-sub | 0.004 | 0.002 |
| HIA | 0.005 | 0.894 |
| F(20%) | 0.003 | 0.021 |
| F(30%) | 0.902 | 0.039 |
| Caco-2 | -4.702 | -5.194 |
| MDCK | 1.44E-05 | 3.05E-05 |
| BBB | 0.134 | 0 |
| PPB | 99.51% | 101.62% |
| VDss | 0.577 | 0.082 |
| Fu | 0.93% | 1.92% |
| CYP1A2-inh | 0.547 | 0.085 |
| CYP1A2-sub | 0.244 | 0.338 |
| CYP2C19-inh | 0.645 | 0.032 |
| CYP2C19-sub | 0.07 | 0.108 |
| CYP2C9-inh | 0.834 | 0.043 |
| CYP2C9-sub | 0.125 | 0.99 |
| CYP2D6-inh | 0.01 | 0.029 |
| CYP2D6-sub | 0.12 | 0.391 |
| CYP3A4-inh | 0.35 | 0.012 |
| CYP3A4-sub | 0.478 | 0.021 |
| CL | 5.204 | 0.723 |
| T12 | 0.716 | 0.026 |
| hERG | 0.188 | 0.008 |
| H-HT | 0.106 | 0.448 |
| DILI | 0.964 | 0.746 |
| Ames | 0.888 | 0.101 |
| ROA | 0.06 | 0.005 |
| FDAMDD | 0.16 | 0.96 |
| SkinSen | 0.766 | 0.444 |
| Carcinogenicity | 0.697 | 0.954 |
| EC | 0.003 | 0.003 |
| EI | 0.063 | 0.866 |
| Respiratory | 0.039 | 0.308 |
| BCF | 1.14 | 0.608 |
| IGC50 | 4.718 | 4.138 |
| LC50 | 5.294 | 5.817 |
| LC50DM | 6.126 | 6.301 |
| NR-AR | 0.107 | 0.001 |
| NR-AR-LBD | 0.088 | 0.004 |
| NR-AhR | 0.948 | 0.733 |
| NR-Aromatase | 0.788 | 0.102 |
| NR-ER | 0.535 | 0.289 |

| compound | C4-31 | Chicago Sky Blue |
|---|---|---|
| NR-ER-LBD | 0.014 | 0.533 |
| NR-PPAR-gamma | 0.844 | 0.002 |
| SR-ARE | 0.81 | 0.496 |
| SR-ATAD5 | 0.784 | 0 |
| SR-HSE | 0.791 | 0 |
| SR-MMP | 0.59 | 0.69 |
| SR-p53 | 0.906 | 0.024 |
| MW | 406.15 | 991.97 |
| Vol | 417.638 | 969.199 |
| Dense | 0.972 | 1.023 |
| nHA | 7 | 22 |
| nHD | 1 | 6 |
| TPSA | 86.22 | 389.2 |
| nRot | 10 | 11 |
| nRing | 3 | 6 |
| MaxRing | 10 | 10 |
| nHet | 7 | 30 |
| fChar | 0 | 0 |
| nRig | 20 | 44 |
| Flex | 0.5 | 0.25 |
| nStereo | 0 | 0 |
| NonGenotoxic_Carcinogenicity | 0 | 0 |
| LD50_oral | 0 | 0 |
| Genotoxic_Carcinogenicity_Mutagenicit | 1 | 12 |
| SureChEMBL | 0 | 3 |
| NonBiodegradable | 0 | 2 |
| Skin_Sensitization | 2 | 5 |
| Acute_Aquatic_Toxicity | 0 | 0 |
| Toxicophores | 3 | 6 |
| QED | 0.351 | 0.041 |
| Synth | 2.057 | 4.636 |
| Fsp3 | 0.174 | 0.059 |
| MCE-18 | 17 | 42 |
| Natural Product-likeness | -1.223 | -0.144 |
| Alarm_NMR | 2 | 9 |
| BMS | 0 | 3 |
| Chelating | 0 | 0 |
| PAINS | 0 | 1 |
| Lipinski | Accepted | Rejected |
| Pfizer | Accepted | Accepted |
| GSK | Rejected | Rejected |
| GoldenTriangle | Accepted | Rejected |

**Supplementary Table 11: Primers for RT-qPCR**

| Primers | Sequence |
|---|---|
| SDHA-F | CAGCATGTGTTACCAAGCTGT |
| SDHA-R | GGTGTCGTAGAAATGCCACCT |
| SDHB-F | ACAGCTCCCCGTATCAAGAAA |
| SDHB-R | GCATGATCTTCGGAAGGTCAA |
| SDHC-F | TTGCTGAGACACGTTGGTCG |
| SDHC-R | GAGACAGAGGACGGTTTGAAC |
| SDHD-F | ATTTCTTCAGGACCGACCTATCC |
| SDHD-R | CAGCCTTGGAGCCAGAATG |
| beta_actin_F | AATCTGGCACCACACCTTCTAC |
| beta_actin_R | ATAGCACAGCCTGGATAGCAAC |
| FLAD1-F | TGACCCCTACTCCTGTAGCC |
| FLAD1-R | AGCTGACGCAGAAAATCCCA |
